# Improving the clinical performance of blood-based DNA methylation biomarkers utilizing locus-specific epigenetic heterogeneity

**DOI:** 10.1101/579839

**Authors:** Brendan F. Miller, Thomas R. Pisanic, Gennady Margolin, Hanna M. Petrykowska, Pornpat Athamanolap, Alexander Goncearenco, Akosua Osei-Tutu, Christina M. Annunziata, Tza-Huei Wang, Laura Elnitski

## Abstract

**Background:** Variation in intercellular methylation patterns can complicate the use of methylation biomarkers for clinical diagnostic applications such as blood-based cancer testing. Here, we describe development and validation of a methylation density binary classification method called EpiClass (available for download at https://github.com/Elnitskilab/EpiClass), that can be used to predict and optimize the performance of methylation biomarkers, particularly in challenging, heterogeneous samples such as liquid biopsies. This approach is based upon leveraging statistical differences in single-molecule sample methylation density distributions to identify ideal thresholds for sample classification.

**Results:** We developed and tested the classifier using reduced representation bisulfite sequencing (RRBS) data derived from ovarian carcinoma tissue DNA and controls. We used these data to perform *in silico* simulations using methylation density profiles from individual epiallelic copies of *ZNF154*, a genomic locus known to be recurrently methylated in numerous cancer types. From these profiles, we predicted the performance of the classifier in liquid biopsies for the detection of epithelial ovarian carcinomas (EOC). *In silico* analysis indicated that EpiClass could be leveraged to better identify cancer-positive liquid biopsy samples by implementing precise thresholds with respect to methylation density profiles derived from circulating cell-free DNA (cfDNA) analysis. These predictions were confirmed experimentally using DREAMing to perform digital methylation density analysis on a cohort of low volume (1-mL) plasma samples obtained from 26 EOC-positive and 41 cancer-free women. EpiClass performance was then validated in an independent cohort of 24 plasma specimens, derived from a longitudinal study of 8 EOC-positive women, and 12 plasma specimens derived from 12 healthy women, respectively, attaining a sensitivity/specificity of 91.7%/100.0%. Direct comparison of CA-125 measurements with EpiClass demonstrated that EpiClass was able to better identify EOC-positive women than standard CA-125 assessment. Finally, we used independent whole genome bisulfite sequencing (WGBS) datasets to demonstrate that EpiClass can also identify other cancer types as well or better than alternative methylation-based classifiers.

**Conclusions:** Our results indicate that assessment of intramolecular methylation density distributions calculated from cfDNA facilitate the use of methylation biomarkers for diagnostic applications. Furthermore, we demonstrated that EpiClass analysis of *ZNF154* methylation was able to outperform CA-125 in the detection of etiologically-diverse ovarian carcinomas, indicating the broad utility of *ZNF154* for use as a biomarker of ovarian cancer.

## Background

A primary aim in cancer diagnostics is to identify and develop biomarkers capable of detecting or assessing malignancies in a reliable and noninvasive manner. Epigenetic alterations have shown significant potential as cancer biomarkers, as many genomic loci become aberrantly methylated during tumorigenesis and can thus serve as indicators of disease (1–3). A particularly attractive application for methylation biomarkers is use in liquid biopsies of peripheral blood, which provide a minimally-invasive means of assessing cancer-specific epigenomic alterations contained in circulating tumor DNA (ctDNA) using a simple blood draw (4).

While genome-wide methylation analysis techniques, such as whole-genome bisulfite sequencing (WGBS) (5) and Infinium BeadArrays (6), have been used to identify scores of differentially-methylated genomic loci in cancer tissues (7), only a handful of methylation biomarkers have been implemented in the clinic (8,9). This is due in part to a number of technical and logistical hurdles involved in translating promising tissue-based methylation biomarkers for use in liquid biopsies, including: (i) the small proportion of plasma ctDNA relative to cell-free DNA (cfDNA) derived from healthy cells (10), (ii) heterogeneity of methylation patterns at a given locus (11–15), (iii) age-associated accrual of methylation (16), (iv) technical artifacts due to bisulfite conversion (17), and (v) differences in the yield of extracted cfDNA between individual or batches of liquid biopsy samples (18). Collectively, these issues can often make it difficult to achieve the high degrees of sensitivity and specificity necessary to attain adequate clinical performance with a discrete number of biomarkers using conventional diagnostic approaches (19). There thus remains an unmet need for the development and implementation of new methods capable of better distinguishing cancer-specific methylation from background methylation “noise” at individual loci in order to harness the diagnostic potential of methylated biomarkers in general.

While assessment of locus-specific differential methylation can be used to readily identify cancer-specific signatures at the tissue-level (20), detection of this differential methylation can become difficult in heterogenous or highly-dilute clinical samples such as liquid biopsies. In these cases, current strategies have primarily focused on identifying and detecting cancer-specific CpG methylation patterns (21–25). Although DNA methylation is regarded as a stable and heritable epigenetic mark, the fidelity and maintenance of specific methylation patterns at the CpG level can vary across the genome (26–29). Differences in methylation levels have been reported between individuals (30), dizygotic twins (31), and within individuals over time (32), suggesting the influence of both genetic and environmental effects on DNA methylation stability and its heritability (33). In the context of cancer, it has been observed that localized methylation patterns become stochastically disordered within individual cells of tumors, creating intratumoral, heterogeneous patterns of methylation (34,35). Importantly, these regions of increased methylation discordance also tend to coincide with those specifically differentially methylated in cancer. Taken together, there is a need for tools to optimize the detection of cancer-specific methylation signatures while accounting for locus-specific heterogeneous populations of methylation patterns both between and within individuals that can complicate the performance of methylation biomarkers (36–38).

The overall aim of the present study was to develop a means of exploiting differences in locus-specific heterogeneous methylation patterns (arising from biological and technical noise but also specifically from tumors) to better distinguish case and control samples for clinical cancer diagnostic applications. In contrast to genome-wide methylation analyses, which are most often employed to identify methylation biomarkers, our approach offers a locus-specific technique for the clinical implementation of methylation biomarkers, particularly in challenging samples. Specifically, we sought to devise an approach to maximize the performance of methylation biomarkers for use with blood-based (liquid biopsy) detection assays. For this purpose, we introduce a binary classification algorithm called *epiallelic methylation classifier* (EpiClass) that can be used to identify and leverage statistical differences in methylation density profiles to improve the overall diagnostic performance of putative methylation biomarkers. As this approach is based on the measurement of methylation density, it is agnostic to methylation pattern permutations, thereby conferring the potential to outperform methods that are reliant on predefined methylation patterns such as methylation-specific PCR (MSP) (39), or calculations of the average methylation level at a locus, which often become ineffective at dilute DNA concentrations. Overall, we reasoned that assessment and analysis of intramolecular methylation densities, i.e. the proportion of CpG sites that are methylated within each epiallelic copy (40), should provide a means of improving both the sensitivity and specificity of methylation biomarkers in clinical diagnostic applications.

We investigated this approach by examining methylation at the *ZNF154* locus, which we previously demonstrated to be differentially-methylated in at least 14 different solid cancer types (2,41) and has additionally been shown by us and others to be a particularly promising methylation biomarker of epithelial ovarian carcinoma (EOC) (8,42). However, the use of *ZNF154* in the context of liquid biopsies is expected to be potentially confounded by the presence of hypermethylated cfDNA derived from healthy tissues of the gastrointestinal tract (43). We therefore reasoned that *ZNF154* would make an ideal model biomarker to test the utility of EpiClass approach.

We first establish the potential utility of EpiClass approach by using RRBS methylation data generated from healthy and cancerous tissues in the framework of *in silico* dilution experiments. This demonstrates the feasibility of improving diagnostic performance through characterization of intramolecular methylation densities in dilute liquid biopsy samples. We then experimentally confirm the predictions of EpiClass performance by generating cfDNA methylation density profiles from liquid-biopsy-derived cfDNA using our previously-reported quasi-digital, high-resolution melt approach called DREAMing to directly assess intramolecular methylation density at single-molecule sensitivity (44). In total, plasma from 34 patients with refractory EOC and 53 cancer-free controls are analyzed as training and testing datasets. From this data we demonstrate significant improvement in the ability to distinguish between patient and control plasma samples using EpiClass compared to calculating the mean locus methylation signal. Furthermore, we show that employment of EpiClass in the assessment of *ZNF154* methylation can more accurately identify EOC in liquid biopsies from etiologically-diverse tumor ovarian cancer subtypes (i.e., both serous and endometrioid tumors) than measurement of blood CA-125, the most commonly-employed biomarker for monitoring EOC. Lastly, using WGBS data generated from plasma of 30 patients with hepatocellular carcinoma and 36 normal controls, we compare the performance of EpiClass to alternative DNA methylation-based analysis methods and demonstrate the ability to apply EpiClass towards assessment of additional cancer biomarkers. Our results suggest that EpiClass offers the potential as an improved screening method for ovarian cancer and more broadly as a practical means of improving the diagnostic performance of DNA methylation-based biomarkers, particularly in challenging samples such as liquid biopsies.

## Results

### Development of a methylation density binary classifier

Figure 1A illustrates the basic principle of the EpiClass approach for classifying samples based on intramolecular methylation density distributions. In this highly simplified example, identification of cancer-specific hypermethylation is obscured by the presence of a few heterogeneously-methylated epialleles in the control sample. In this situation, not uncommon in cfDNA methylation analyses, the samples cannot be differentiated by assessment of mean methylation (or β-values) at individual CpGs, the entire locus or even only fully-methylated epialleles (Supplementary Figure S1). In contrast, by considering the methylation density of individual cfDNA fragments, or epialleles, and their relative abundances or fractions in each sample, i.e. the epiallelic fractions, the case is distinguishable from the control sample (Figure 1B).

**Figure 1:**
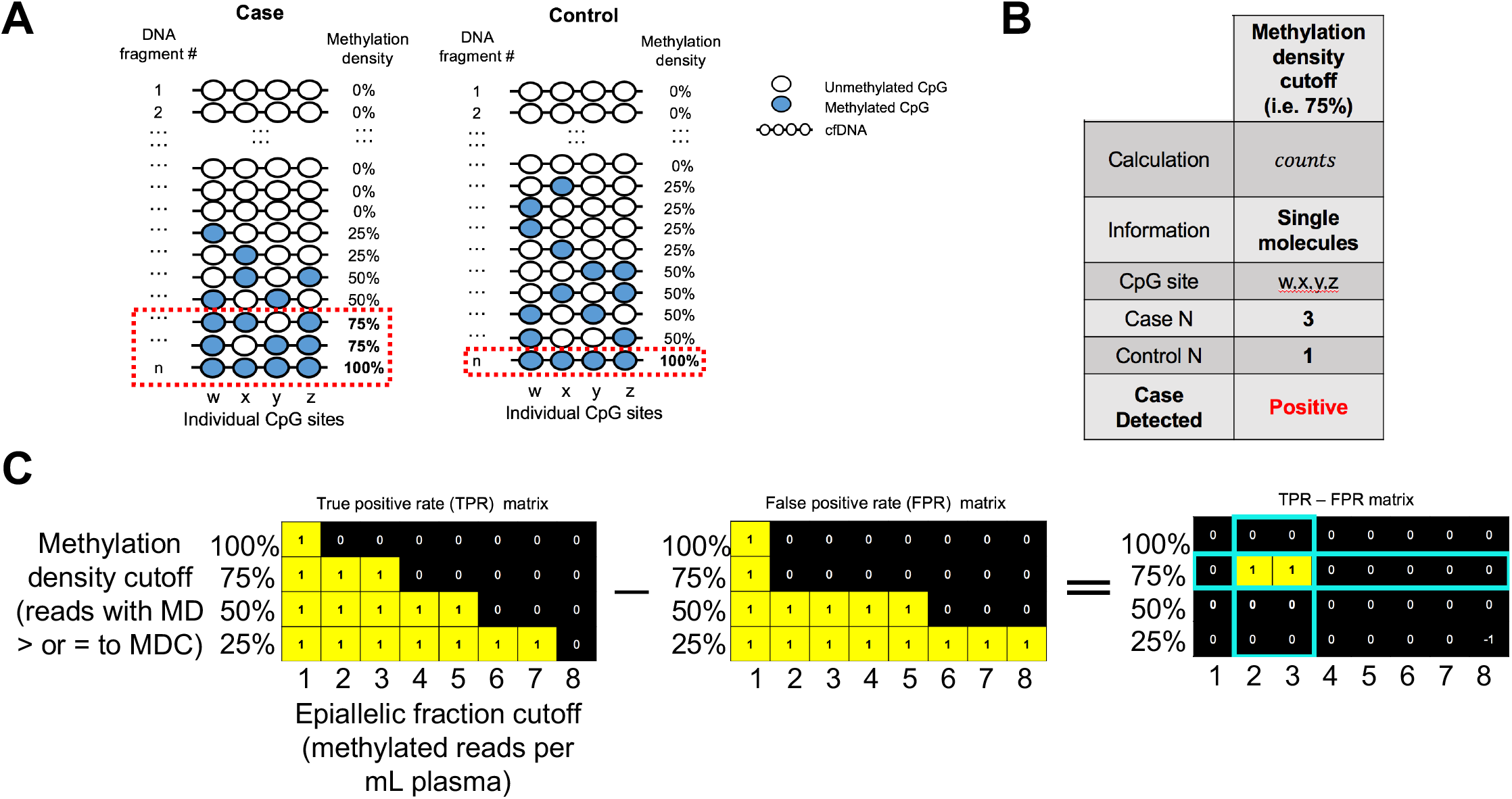
Schematic of the methylation density binary classifier. **A**) Example illustration of heterogeneous mixtures of cfDNA epialleles with different methylation densities in case and control liquid biopsy samples. **B**) Table characterizing the use of a methylation density classifier cutoff. **C**) Schematic illustrating the EpiClass procedure for determining optimal methylation density and epiallelic fraction cutoffs.

In order to further leverage the discriminatory power of using methylation-density-determined epiallelic fraction, we sought to codify this approach by creating a streamlined, biostatistical tool called *epiallelic methylation classifier* (EpiClass). Implementation of EpiClass requires a dataset that provides a representative distribution of epiallelic methylation densities at the locus of interest, such as can be derived from standard bisulfite sequencing data (WGBS, RRBS, and other) methods. Notably, positional information of CpG sites are not required, thereby making EpiClass compatible with digital or quasi-digital high-resolution DNA melting techniques, such as HYPER-Melt or DREAMing techniques (44,45), respectively. The algorithmic procedure for EpiClass is shown in Figure 1C, which is used to identify the methylation density and epiallelic fraction cutoffs that optimize classification of case and control samples. Methylation of a finite number of CpG sites in the analyzed locus results in a tabularized distribution of methylation densities in case and control sample sets. This table is then used to calculate the true positive and false positive rates as determined by iteratively varying methylation density and epiallelic fraction cutoffs. Figure 1C shows the TPR and FPR matrices generated for the example scenario of Figure 1A, with the corresponding optimal methylation density and epiallelic fraction cutoffs found by taking the cutoff combination that maximizes the positive difference between the TPR and FPR (in this simplified case, TPR is 100% and FPR is 0%). These cutoffs are the predicted optimal cutoffs for the genomic locus in the test sample set, which can then be further validated in a secondary cohort of samples.

### Simulated dilute admixtures for predicting EpiClass performance

We previously reported the presence of aberrant methylation at the *ZNF154* locus in at least 14 different solid cancer types (2,41) and also established the locus as one of the most promising candidate biomarkers for epithelial ovarian carcinoma (EOC) (8). However, analysis of Illumina Infinium Human Methylation 450K array data from the Cancer Genome Atlas (TCGA) (46) revealed that healthy colorectal tissues also exhibit hypermethylation at this locus (Supplementary Figure S2), portending limitations in the utility of *ZNF154* as a biomarker in liquid biopsies derived from peripheral blood. Additional analyses of RRBS data from EOCs, healthy ovarian tissues, and peripheral blood samples from healthy patients also revealed heterogeneous methylation at this locus across individual reads and samples that might further complicate analyses (Supplementary Figure S3). Therefore, in spite of the promising discriminatory potential of hypermethylation at *ZNF154* to classify ovarian carcinomas (41), methylation heterogeneity at this locus would be expected to challenge its utility as useful liquid biopsy marker, thereby making this locus an ideal candidate for testing the EpiClass approach.

In order to further validate the selection of *ZNF154* for our studies, we sought to develop a generalizable approach for predicting whether implementation of EpiClass with a given methylation biomarker could improve diagnostic performance over other methods in liquid biopsy specimens. Furthermore, we reasoned that such an approach might also aid in the selection of biomarkers that could benefit from the EpiClass method in general. Toward this end, we investigated whether methylation density distributions derived from publicly-available datasets might be used to create simulated dilute admixtures resembling cfDNA solutions. Simulated cfDNA solutions were created using *in silico* admixtures from RRBS reads of EOCs (n=12) and white blood cells (WBCs; n=22). In particular, cancer-positive cfDNA samples were simulated using a bootstrapping method by combining randomly-sampled RRBS reads from each of the EOC samples after being randomly-paired and mixed with a WBC background sample at various EOC:WBC admixture ratios ranging from 100% down to 0.01%. These were then compared with RRBS reads generated exclusively from the WBCs. Data from this evaluation were used to identify methylation density cutoff values achieving the highest area under the receiver operating characteristic curve (AUC) value based on EpiClass analysis, as determined by sample sets at each admixture ratio. The AUC values achieved by implementation of EpiClass to classify the samples were compared to those achieved using overall mean locus methylation as a baseline classification approach.

Figure 2 shows the AUC performance of EpiClass versus average methylation values over the entire range of simulated EOC:WBC admixture ratios. The results of this analysis indicate that EpiClass classification outperformed average methylation classification at all admixture ratios, particularly as the EOC:WBC ratio decreased below the 1% abundance commonly reported for ctDNA in cfDNA from cancer patients (47). We then generated a second simulated cohort of 50 cancer-positive and 50 cancer-free cfDNA samples, to observe the variance in EpiClass performance in simulated cfDNA solutions of 1%, 0.1% and 0.01% EOC:WBC admixture ratios. The results of these simulations, shown in Supplementary Figure S4 and Supplementary Figure S5, indicate that the probability of improving sample classification relative to mean locus methylation steadily increases as the methylation density cutoff increases from 20% to approximately 85% at all admixture ratios. Performance drops off as the admixture ratio approaches 0.01% and, interestingly, at methylation density cutoffs >85%, as might be observed in traditional assays such as MSP due to stochastic sampling of only heavily- or fully-methylated epialleles.

**Figure 2:**
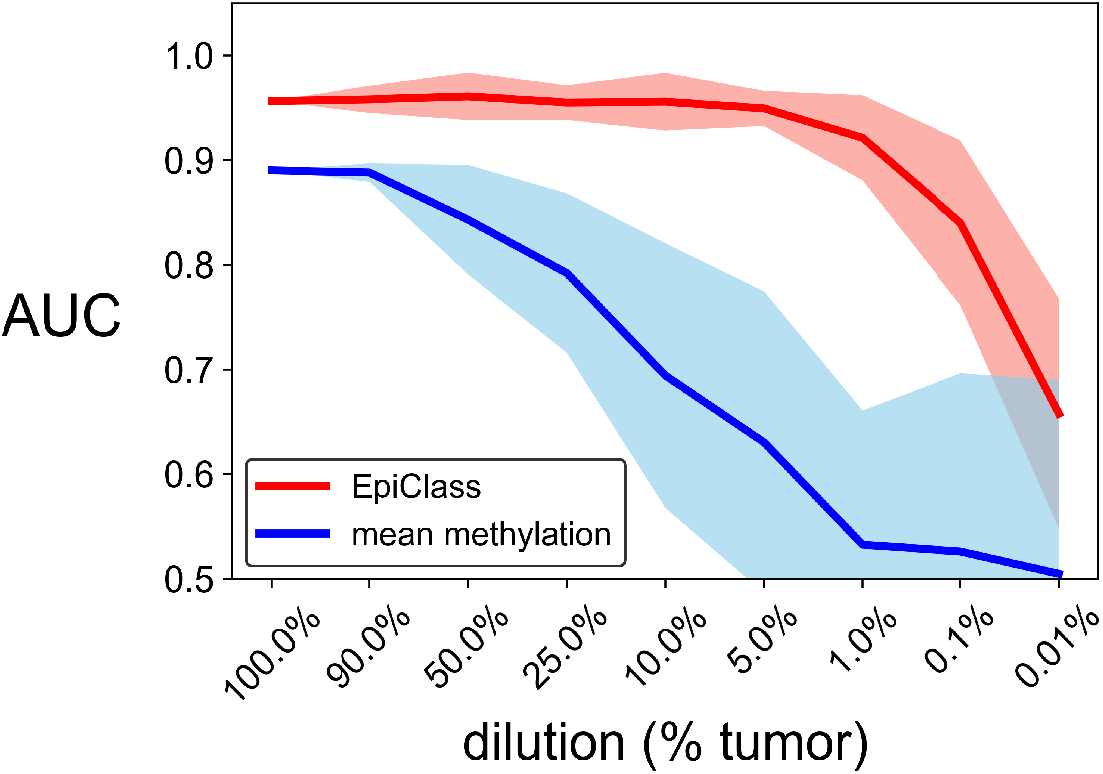
Simulated performance of EpiClass as a function of admixture ratios of ovarian carcinoma (EOC) to WBC RRBS reads. The performance of the methylation density binary classifier (EpiClass, *red*) and mean locus methylation classifier (*blue*) at increasing dilutions of EOC RRBS reads in a background of WBC RRBS reads acquired from Widschwendter *et al* (8). Simulated performance of improved classification over mean methylation at different read depths demonstrated in Supplementary Figure S4 and S5. Abbreviations: AUC = area under the curve; EpiClass = methylation density classifier.

### Implementation of EpiClass in liquid biopsies

We next validated the predicted performance of EpiClass using primary data derived from a cohort of liquid biopsies from 67 patients (26 women with late-stage ovarian cancer and 41 cancer-free women; Supplementary Table S1). For these samples, we obtained methylation densities of individual DNA fragments using DREAMing melt profiles (44) (see Supplementary Figure S6 for examples). We further validated the methylation densities obtained by DREAMing with bisulfite sequencing. The two methods showed good agreement (Supplementary Figure S7) indicating sufficient congruency to implement DREAMing as a simpler and cheaper method for locus-specific methylation density analysis than deep bisulfite sequencing.

As shown in Figure 3A and Supplementary Figure S8, preliminary meta-analysis of the methylation density profiles of EOC-positive and cancer-free women revealed that while both cohorts exhibited a large fraction of *ZNF154* epialleles with little to no methylation, only cfDNA from cancer-positive patients exhibited the presence of a significant proportion of densely-methylated *ZNF154* epialleles. However, comparison of the overall population of epialleles indicated that there was no significant difference in mean methylation between cases and controls, suggesting that classification by mean methylation was not sufficient to discriminate samples (Supplementary Figure S9).

**Figure 3:**
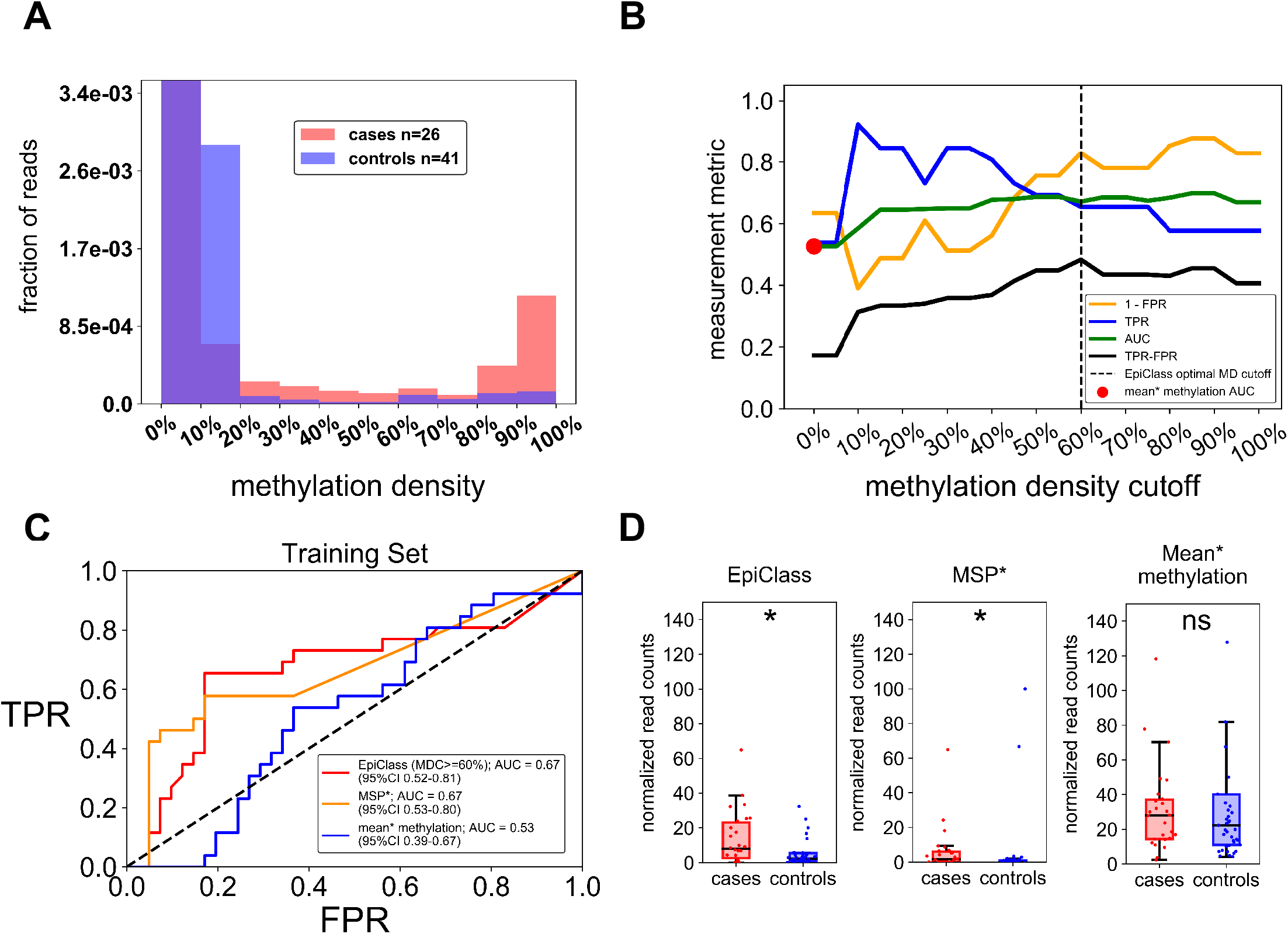
Performance of EpiClass in EOC patient and control plasma samples. **A**) The pooled epiallelic fractions of cfDNA methylated epialleles with varying methylation densities in EOC (*red*, n=26) and healthy (*blue*, n=41) patient plasma samples. Purple-shaded regions indicate overlap between the two plasma sets. **B**) Performance of EpiClass at each methylation density cutoff for the EOC and healthy control plasma samples. Dotted line shows the optimal methylation density cutoff derived from EpiClass. The red dot indicates the ROC curve AUC for the mean methylation cutoff. Measurement metric refers to either 1 – FPR, TPR, AUC, or TPR – FPR. **C**) ROC curves showing the classification performance of using the optimal methylation density cutoff determined by EpiClass (*red*), MSP (*orange*), or mean methylation cutoff (*blue*) to identify the EOC and healthy control plasma samples. **D**) Boxplots showing the performance of the epiallelic fraction cutoffs for either the optimal 60% methylation density cutoff determined by EpiClass, MSP, or mean methylation to classify plasma samples from EOC patients (*red*, n=26) or healthy controls (*blue*, n=41) Y-axes adjusted to ignore healthy control outliers. Abbreviations: EOC = epithelial ovarian carcinoma; EpiClass = methylation density classifier; MDC = methylation density cutoff; AUC = area under the curve; * indicates p < 0.05, two-sided Wilcoxon rank-sum test; ns = not significant; Mean* methylation was inferred from the fraction of all methylated epialleles. MSP* performance estimated using an MDC of 95%. Supplementary Figure S9 demonstrates no statistical difference between sample cohorts with respect to mean methylation.

We next employed EpiClass to identify methylation density and epiallelic fraction cutoffs that would optimize the diagnostic performance for classifying EOC-positive women based solely on their respective liquid biopsy *ZNF154* methylation density profiles. The results of EpiClass analysis, shown in Supplementary Figure 10, identified a methylation density cutoff of 60% and a normalized epiallelic fraction of 6.7 epialleles per mL of plasma (viz., 6.7 epialleles, at least 60% methylated) as the optimal cutoff values for maximizing diagnostic performance. Figure 3B shows the classification performance as a function of methylation density cutoff (using corresponding optimal epiallelic fraction cutoffs) in comparison to classification by mean methylation alone. Data from this analysis demonstrate that overall performance increases as the methylation density cutoff is increased to 20%, at which point it remains largely flat, agreeing with the predicted performance of the previous simulation. This result indicates that consideration of heterogeneously-methylated epialleles can significantly improve classification performance. In contrast, consideration of epialleles at methylation densities below 20% reduces overall performance (due to high fractions of epialleles with low methylation density in the controls) by effectively reducing clinical specificity. The optimal methylation density cutoff, defined as the maximum positive difference between TPR and FPR, was identified here as 60%. These points are further illustrated in Figure 3C-D, which shows that EpiClass achieves better performance than using a 0% (mean methylation) methylation density cutoff (EpiClass AUC = 0.67, optimal sensitivity/specificity = 65%/83% vs. mean methylation AUC = 0.53, optimal sensitivity/specificity = 54%/63%). Consideration of only heavily methylated epialleles, as targeted in MSP-based techniques, was defined by epialleles with 95% or more methylation density and yielded the same AUC and specificity as EpiClass (AUCs: 0.67 vs 0.67; specificities: 83% vs 83%), however the EpiClass optimal sensitivity was higher (65% vs 58%), suggesting heterogeneously-methylated epialleles allow for a modest improvement in detection. The failure of the mean methylation level to classify any cases is likely a result of low-level background methylation noise contributed by the higher abundance of low-methylation-density epialleles in the control samples than the cases.

### Independent validation of EpiClass threshold values

We validated the methylation density and epiallelic fraction cutoffs identified by EpiClass in our initial cohort by applying them to the analysis of a second, independent cohort. For this validation cohort, we used archived plasma samples (n=36), comprising 3 separate blood draws obtained from 8 women over a longitudinal study period of 8 weeks (n=24), as well as plasma samples obtained from cancer-free women (n=12; Supplementary Table S2). Initial analysis, shown in Figure 4A, indicated that these samples exhibited a similar overall methylation density profile as the first cohort in that the cases had higher levels of heterogeneously-methylated epialleles than in the control cohort samples, many of which also exhibited the presence of hypermethylated epiallelic variants.

**Figure 4:**
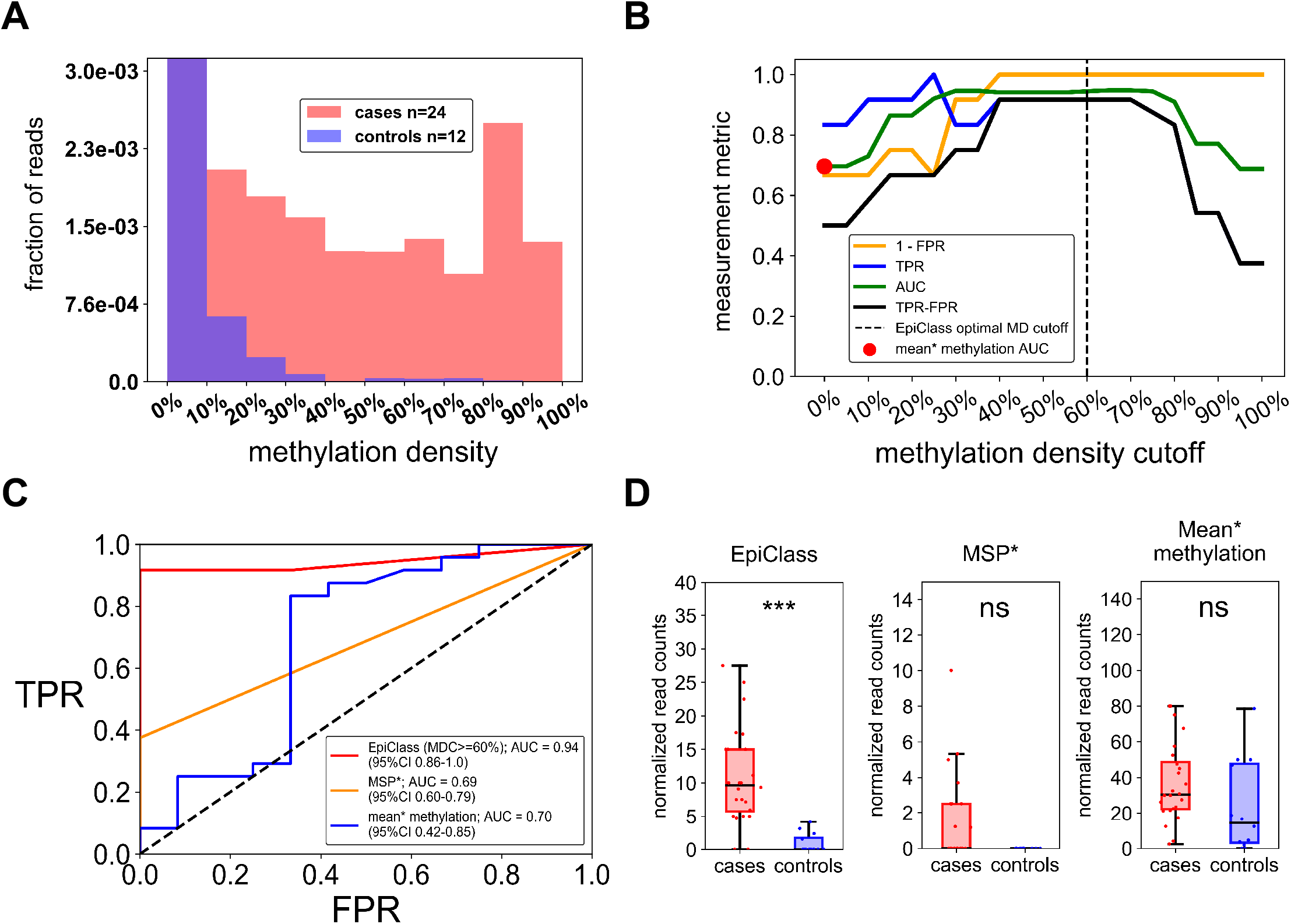
Validation of EpiClass cutoff values and corresponding performance for identifying EOC from patient plasma. **A**) The pooled epiallelic fractions of cfDNA methylated epialleles with varying methylation densities in the second EOC (*red*, n=24) and healthy (*blue*, n=12) patient plasma sample cohort. Purple shaded regions indicate overlap between the two plasma sets. **B**) Performance of EpiClass at each methylation density cutoff for the EOC and healthy control plasma samples. Dotted line shows the optimal methylation density cutoff derived from EpiClass. The red dot indicates the ROC curve AUC for the mean methylation cutoff. Measurement metric refers to either 1 – FPR, TPR, AUC, or TPR – FPR. **C**) Receiver operating characteristic curve for the optimal 60% methylation density cutoff on the second plasma cohort. **D**) Boxplots indicating the distribution of sample normalized read counts with intramolecular methylation densities greater than or equal to 60% (EpiClass), 95% (MSP*), or greater than 0% (mean* methylation). Abbreviations: EOC = epithelial ovarian carcinoma; EpiClass = methylation density classifier; MDC = methylation density cutoff; AUC = area under the curve; *** indicates p < 0.001, two-sided Wilcoxon rank-sum test; ns = not significant; Mean* methylation was inferred from the fraction of all methylated epialleles. MSP* performance estimated using an MDC of 95%.

Independent evaluation of the methylation density cutoffs from the validation cohort, as shown in Supplementary Figure S11 and Figure 4B, demonstrated that the ideal methylation density cutoff values identified by EpiClass were largely consistent between the simulated (20-85%), training (20-95% density) and validation (20-80% density) cohorts. These results indicate that experimentally-determined methylation density thresholds are likely consistent and applicable when evaluating independent cohorts with similar patient characteristics. Figure 4C and 4D show that classification of the validation set using the optimal methylation density cutoff of 60% identified in the training set achieved 91.7%/100.0% sensitivity and specificity, in which the sensitivity plus specificity was higher than either the mean methylation level (83.3%/66.7% sensitivity and specificity) or the 95% methylation density cutoff (37.5%/100.0% sensitivity and specificity). Again, mean methylation suffered from a loss of clinical specificity likely due to heterogeneous, low-methylation-density epialleles whereas consideration of heavily-methylated epialleles compromised clinical sensitivity as not all case samples possessed significant fractions of heavily-methylated epialleles.

### Performance of EpiClass vs CA-125 in liquid biopsies from patients with EOC

Next, we blindly classified these samples according to the 60% optimal methylation density and 6.7 epialleles per mL plasma epiallelic fraction cutoffs established by EpiClass analysis of the training cohort, the results of which are shown in Figure 5. We compared the classification performance using the EpiClass-identified methylation density cutoffs with gold-standard patient CA-125 levels (available for all but one of the samples in the validation set). Figure 5 shows that EpiClass correctly classified 16 of 23 (69.6%) samples, whereas CA-125 correctly classified only 11 of 23 (47.8%) using the standard clinical CA-125 cutoff of 35 U/mL. Of additional note, CA-125 misclassified all of the patients with non-serous ovarian cancer subtypes (n=9), whereas EpiClass-thresholded *ZNF154* correctly classified 6 of the 9 endometrioid samples. This finding suggests that EpiClass analysis of *ZNF154* may represent a viable alternative to CA-125 for companion diagnostics of non-serous ovarian cancer. Combining CA-125 measurements with EpiClass cutoffs improved classification performance even further, achieving correct classification of 6 of 9 non-serous and 14 of 14 serous for a total of 20 of 23 (87.0%) cases. Here, too, implementation of EpiClass-derived cutoffs for *ZNF154* methylation achieved 100.0% clinical specificity.

**Figure 5:**
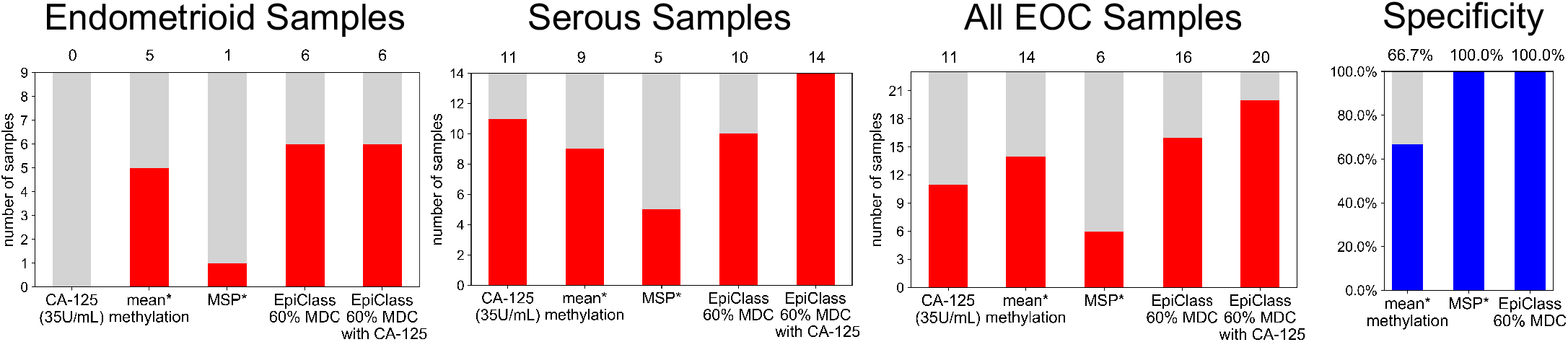
Counts of endometrioid (n=9) or serous (n=14) EOC subtype patient plasma samples from the validation cohort above the CA-125 cutoff (35 U/mL) or EpiClass cutoffs derived from the training cohort. Numbers above each bar indicate the number of samples above each given classification cutoff. Mean* methylation was inferred from the fraction of all methylated epialleles. MSP* performance estimated using an MDC of 95%. No available specificity measurements for CA-125 as the control samples (n=12) did not have measured CA-125 U/mL concentrations. Abbreviations: MDC = methylation density cutoff.

### Comparison of EpiClass to alternative methylation-based sequencing analyses

We compared the classification performance of EpiClass to another cfDNA methylation analysis tool, CancerDetector (25), which has been previously reported to provide higher liquid biopsy cancer detection performance than other epigenetic diagnostic methods, including CancerLocator (48) and methylation haplotype blocks (21). For this comparison, we used whole genome bisulfite sequencing (WGBS) data of cfDNA derived from liquid biopsies obtained from 30 patients diagnosed with hepatocellular carcinoma (HCC) (from Chan *et. al*. (24), EGAS00001000566, and Li *et. al*. (25), EGAD00001004317), which included 25 patients with early-stage disease (24), and 36 normal controls. Using this data, we assessed methylation at our locus of interest, *ZNF154,* and three additional liver cancer loci, using both methods. The samples were randomly split into 10 training and test sample sets. Training samples from each run were used to construct classifiers based on methylation density cutoffs (for EpiClass) or Tumor and Normal Class beta distribution shape parameters (in the case of CancerDetector). These classifiers were then applied to the test set samples and the performances of the two methods were compared (see Methods).

Figure 6A and B show that EpiClass slightly outperformed CancerDetector with respect to classifying the WGBS cases and normal plasma samples when utilizing *ZNF154* as the biomarker (EpiClass mean AUC = 0.77 versus CancerDetector mean AUC = 0.70), although this difference was not statistically significant (p = 0.059; Wilcoxon rank sum 2-sided test). For all 10 EpiClass training sample set runs, the ideal methylation density cutoff was determined to be 45% (Supplementary Table S3), which is consistent with the previously determined range of ideal methylation density cutoffs for the EOC training and independent validation cohorts. This further confirms the utility of leveraging heterogeneous methylation data to improve the performance of *ZNF154* and demonstrates that EpiClass derived optimal cutoffs for a given locus are consistent not only across different sample cohorts but also across different cancer types and stages of disease.

**Figure 6:**
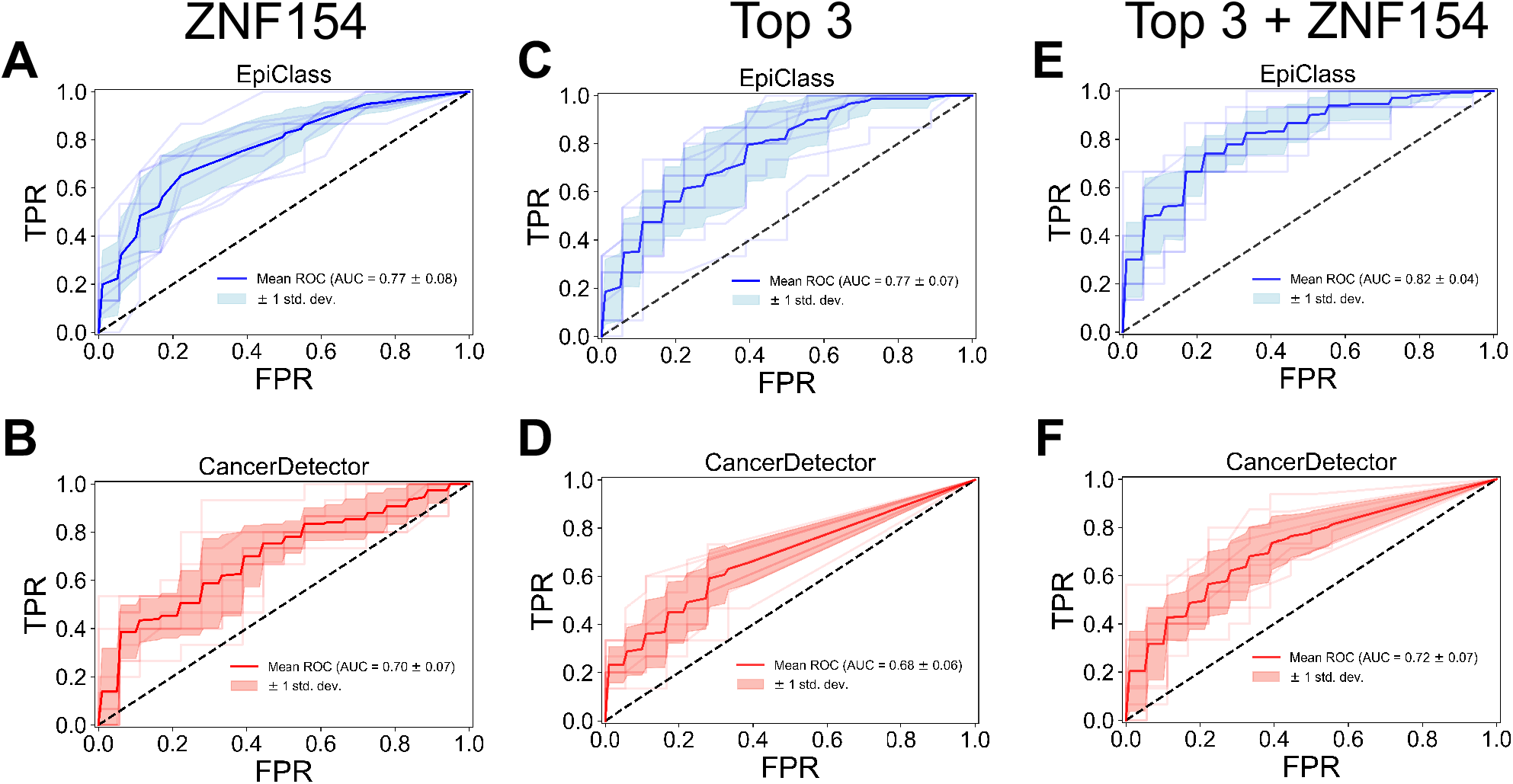
Comparison of EpiClass and CancerDetector. **A-B**) Receiver operating characteristic curves (ROCs) based on classification of 10 test sample sets using either the optimal methylation density read counts at the *ZNF154* marker region for EpiClass (A, *blue*), or the estimated sample tumor fraction derived by CancerDetector (B, *red*) using reads at the *ZNF154* marker region. Light shaded lines indicate individual ROC curves for each test sample set. Dark line indicates the mean ROC curve. Light shaded region indicates 1 standard deviation from the mean. **C-D**) Same as A-B except for the chosen top 3 liver cancer markers. **E-F**) Same as C-D except for the top 3 liver cancer marker regions and *ZNF154* marker region. Abbreviations: AUC = Area under the curve; std. dev. = standard deviation.

To explore the generalizability of EpiClass for use with panels of methylated biomarkers, we identified additional genomic regions that are hypermethylated in TCGA liver hepatocellular carcinoma (LIHC) tumors with respect to normal controls and classified the WGBS samples based on their methylation status at these loci. Specifically, we selected the top three CpG probes with the largest positive median difference in beta-value methylation level between matched tumor and normal samples using LIHC 450K methylation array datasets collected from TCGA. We also implemented a previously-described bioinformatic pipeline to identify highly cancer-specific methylation by only considering probes exhibiting β < 0.2 in all of the normal samples. Additionally, in order to compare the performance of EpiClass to CancerDetector, we limited our probe selection to the liver-cancer-specific markers previously identified by CancerDetector (25). These markers are notable in that they exhibit differential methylation, not only between liver cancer samples and matched normal tissue, but also between liver cancer samples and normal plasma samples. For EpiClass, the methylated status at each of the three loci for the WGBS samples was pooled by constructing a support vector machine (SVM) classifier with a linear kernel and using the normalized methylated read counts at each locus (see Methods). Figure 6C and D show that EpiClass significantly outperforms CancerDetector with respect to classification based on the three loci, (EpiClass mean AUC = 0.77 versus CancerDetector mean AUC = 0.68; p=0.013, Wilcoxon rank sum 2-sided test). In Figure 6E and F, we further expanded the biomarker panel by also including *ZNF154* (4 biomarkers total), resulting in mean AUCs of 0.82 and 0.72 for EpiClass versus CancerDetector, respectively (p=0.005; Wilcoxon rank sum 2-sided test). Overall, EpiClass exhibited better classification performance than CancerDetector in all tested methylation biomarkers.

## Discussion

Recent analyses of cancer methylomes have shown that while cancer-specific hypermethylation can be highly deterministic, methylation patterns between tumor subpopulations at these loci often exhibit considerable epigenetic polymorphisms (13). Diagnostic methylation assays that rely on heavily methylated signatures, such as MSP, may struggle at detecting early stage tumors in part due to their inability to discriminate the subtle changes in methylation density distributions that are likely present in heterogeneous tumors and at early stages of carcinogenesis. In the present work, we report a method for leveraging disease-associated differences in epiallelic methylation density profiles to improve performance of methylation biomarkers for use in clinical diagnostic assays, particularly when evaluating challenging samples such as liquid biopsies. Our results demonstrate that assessment of methylation density information at the level of individual DNA fragments can be used to establish effective thresholds potentially capable of overcoming the inherent biological noise associated with methylation from background sources such as clinical, technical and biological variation or age-related epigenetic drift (49).

In this study, we primarily focused on the *ZNF154* locus based upon previous reports by us and others showing it to be a promising, recurrently-methylated cancer biomarker whose implementation in liquid biopsy is nonetheless complicated by the presence of background methylation from cfDNA derived from healthy tissues (8,41,50). Furthermore, the potential utility of *ZNF154* as a pan-cancer methylation biomarker also afforded the opportunity to directly compare the performance of EpiClass in different cancer types. Our results suggest that EpiClass can be successfully implemented with *ZNF154* cfDNA methylation data obtained by different methods (RRBS, WGBS, DNA melting data via DREAMing) to improve liquid-biopsy-based detection of disparate cancer phenotypes. Likewise, *ZNF154* methylation appears to be useful for detection and monitoring of EOC subtypes, in contrast to CA-125, which is largely limited to only serous-type ovarian carcinomas. Beyond EOC, EpiClass is easily generalizable to individual or multiple cancer biomarkers irrespective of their cancer specificity as long as the appropriate case and control datasets are used.

To demonstrate the generalizability of EpiClass, we also implemented it with independent datasets and methylation biomarkers in the context of HCC. With this data we also compared the classification performance of EpiClass with a previously-described methylation classification tool, CancerDetector. Overall, our results indicate that not only is EpiClass readily generalizable to individual or multiple cancer biomarkers, but that it also can achieve higher performance than other methods applied to the same targets and datasets. Notably, applying EpiClass to *ZNF154* methylation in the context of WGBS data from HCC patient and normal control plasma samples demonstrated promise for detecting early stage hepatocellular carcinoma as 25 out of the 30 WGBS datasets were collected from patients with early stage disease (24). We conclude that EpiClass is generalizable to individual or multiple biomarker panels, applicable to sequencing in addition to DNA melting data, performs as well as CancerDetector for classification of cancer-positive liquid biopsy samples, and that *ZNF154* may also be an effective biomarker for detection of diverse cancer types, including liver and ovarian cancers.

In addition to its utility for predicting and improving the performance of novel methylated biomarkers, EpiClass also has the potential to prompt reevaluation of promising methylation biomarkers that may have been overlooked or excluded due to perceived background noise resulting from heterogeneous methylation. As this noise depends not only on the locus in question but also on the cohort of samples analyzed, EpiClass could be used to prescreen for methylated loci that would have the highest performance specific to the clinical context of interest. For example, in this study, we selected three loci that exhibited liver-cancer-specific differential methylation based on public TCGA data and assessed their classification performance using WGBS data from liquid biopsies obtained from patients diagnosed with HCC and healthy individuals. Interestingly, the performance of these three loci was equivalent to that of using only *ZNF154* in the case of EpiClass analysis, or slightly reduced in the case of CancerDetecter. Consideration of all four loci provided only a minor improvement in analytical performance. In terms of marker screening, *ZNF154* may be a more efficient choice for inclusion in a biomarker panel, whereas the other three loci may be useful for identifying tissue of origin.

It is also possible to the results of EpiClass analyses could be used to inform the most suitable assay method for a candidate methylation biomarker. For example, a locus of interest with a high methylation density optimal cutoff would ostensibly imply high levels of background methylation would likely obfuscate mean methylation analysis but might thereby be well-suited to MSP-based assays targeting heavily methylated epialleles. Alternatively, optimal methylation density cutoffs in the moderate range would imply the presence of heterogeneously-methylated epialleles from healthy tissue(s) and indicate that methods capable of analyzing intramolecular methylation density, such as DREAMing or deep sequencing, may be warranted in challenging samples such as liquid biopsies. A low-level methylation density cutoff would imply an ideal biomarker with minimal background methylation, for which methods sensitive to any level of hypermethylation, e.g. methylation-sensitive high resolution melt (MS-HRM), might be employed (51,52). In theory, EpiClass thresholds might also be adjusted to differentiate even marginal changes in methylation density to increase early stage sensitivity while maintaining an acceptable level of specificity. Future studies will be needed to further explore the relationship between methylation density and the development and progression of cancer.

We observed that the optimal methylation density cutoffs identified by EpiClass were relatively consistent in the present study, however it is probable that the epiallelic fraction cutoff might vary between distinct datasets or analysis techniques as seen by comparison of the training and validation sample sets as well as between tissue and plasma sample sets. This likely reflects inherent differences in the composition of the tumor tissues and suggests that care should be taken when employing EpiClass thresholds in disparate sample types assessed by different or incongruent technologies. Nonetheless, our results indicate that consideration of heterogeneously-methylated epialleles is expected to improve diagnostic performance in various sample types by increasing the separation between tumor and normal signals regardless of sample type or cohort. Additionally, the agreement of the methylation density cutoffs between our different sample cohorts implies that localized discordant methylation profiles are possibly a product of consistent underlying biological phenomena as observed here in the case of EOC and liver cancer at the *ZNF154* locus.

It should be noted that the EpiClass approach is dependent upon sensitive and accurate assessment of methylation density at the individual epiallele level. This can pose technical or logistic barriers, particularly for sequencing-based approaches, which can require significant time and cost to achieve adequate sensitivity and statistical power to determine accurate methylation density profiles in samples containing dilute tumor DNA, such as liquid biopsies. In contrast to methylation patterns, epiallelic methylation density information can be more easily assessed using HRM techniques, such as DREAMing, thereby opening the door for biomarkers to be evaluated via low-cost, nonsequencing-based assays amenable to low-resource clinic settings (44,51). The DREAMing-based technique we employed here offers several advantages for targeted profiling of methylation density. In particular, the short turnaround time for DREAMing (results in several hours) and low cost (approximately $10.00 per sample) make it a practical option for profiling methylation density. Secondly, single molecule sensitivity is readily achievable by DREAMing, which is particularly important when working, as here, with limited (plasma) sample volumes. Furthermore, unlike bisulfite sequencing, DREAMing does not require analysis of sequencing results or patterns, drastically reducing the turnaround time for determining optimal thresholds for maximizing performance or applying established thresholds in a given clinical application.

Unlike other methods that rely upon thousands of genomic loci to classify a sample as cancer-positive, EpiClass is instead designed for the assessment of individual genomic regions. However, as demonstrated in this study, it is also possible to combine counts of methylated fragments from different loci into a learned model after assessing each separately with EpiClass. One reason for assessing hundreds to thousands of loci is to increase the probability of detecting tumor derived DNA with any of the loci, especially when using low coverage sequencing information. Conversely, EpiClass is ideally suited for use with a small panel of loci using high depth data, such as derived from DREAMing, to achieve high classification performance. With low coverage data, relative performance decreases, as expected and as demonstrated here, when comparing DREAMing data versus WGBS data. However, the relatively simple procedure of including heterogeneous methylation as a measurable signal performs as well if not better than methods which rely on methylation patterns or attempt to identify tumor derived reads based on joint methylation probabilities across adjacent CpG sites. Because the majority of methylation patterns in the genome are characterized by bimodality (either heavily methylated or unmethylated) it is reasonable for many classification methods to construct a model that reflects these underlying distributions. However, it can be difficult to assign DNA fragments with heterogeneous methylation patterns to one mode or the other. In liquid biopsies, the dearth of tumor DNA means it is essential to capture as much signal as possible. The ability to include heterogeneously-methylated tumor-derived DNA using a simple threshold is one way to increase signal and improve detection sensitivity. Applying this method to a small panel of biologically relevant loci at high depth may greatly improve the ability to detect disease.

## Conclusions

The EpiClass approach presented here is a relatively simple, effective, and interpretable technique for overcoming issues related to the presence of heterogeneous methylation patterns that arise from inherently stochastic cellular processes associated with the accumulation and maintenance of DNA methylation, such as localized discordant methylation reflective of intratumor heterogeneity. Likewise, this approach is likely to be largely suitable not only for optimizing the performance of methylation biomarkers for clinical diagnostic applications such as liquid biopsies, but also more generally for the study and evaluation of any biological phenomena associated with the dynamic localized accumulation or loss of DNA methylation, particularly in complex, composite or dilute samples.

## Methods

### Datasets and samples

#### 450K Illumina Infinium HumanMethylation450 BeadChip datasets

Processed Illumina Human 450K data for EOCs (GEO accession GSE72021, described in (8,53)) from 221 tumor samples (171 serous, 18 endometrioid, 14 clear cell, 9 mucinous and 9 other histological cancer subtypes) and WBCs (GEO accession GSE55763, described in (54)) from 2,664 individuals was downloaded from the NCBI’s Gene Expression Omnibus (GEO, https://www.ncbi.nlm.nih.gov/geo/). Data for TCGA solid tumor and control sample sets was downloaded from the Broad Institute (https://gdac.broadinstitute.org/) FireHose. Beta-values for probes cg11294513, cg05661282, cg21790626, and cg27049766 were extracted using custom Python-2.7.14 (https://python.org) scripts.

#### Reduced representation bisulfite sequencing data

Reduced representation bisulfite sequencing (RRBS) data was obtained for 12 EOCs, 10 healthy ovarian tissues, and 22 WBC samples used in Widschwendter *et al.* (8). The data was downloaded from the European Genome-phenome Archive (dataset accession: EGAD00001003822).

#### Plasma samples

All samples for this study were obtained after approval by institutional review board (IRB) and the study was conducted in accordance with the U.S. Common Rule. Plasma samples (n=107) were obtained from 34 patients with late-stage residual EOC and 57 pathologically normal control patients with written consent given by (all) the participant subjects. The patients with EOC were derived from two different clinical trials, described below, and controls were recruited by Fox Chase Cancer Center. Samples were split into two cohorts. One was used to assess the performance of EpiClass and establish optimal cutoffs in plasma and the second cohort was used to validate EpiClass cutoffs in plasma and compare the cutoffs to CA-125.

The first cfDNA cohort contained plasma from 26 EOC-positive patients and 41 healthy women (1 sample per patient). The patients with EOC had previously received standard-of-care treatment (a platinum- and taxane-containing regimen), relapsed after one or more subsequent treatment regimens, and been recruited into a clinical trial testing combined treatment with bevacizumab and dasatinib (NCT01445509). The plasma samples were taken at baseline before treatment was administered. Volumes of plasma processed and extracted cfDNA concentrations can be found in Supplementary Table S1.

The second cohort encompassed 24 plasma samples from 8 patients with ovarian cancer (3 samples per patient) and 12 control samples from healthy women (1 sample per patient). Here, too, patients with ovarian cancer were given standard of care, relapsed, and were recruited to a clinical trial, this one testing combined treatment with bevacizumab and sorafenib (NCT00436215) (55). Ovarian cancer patient plasma samples were taken at three separate time points over 6 weeks (at baseline, 2 weeks after exposure to the first agent, and 2 weeks after exposure to the second agent). Blood was collected into standard EDTA tubes and maintained on ice for transport; plasma was separated immediately using centrifugation and frozen in 1.0- to 1.5-ml aliquots at −80°C. Additionally, CA-125 protein levels in the blood were measured every 4 weeks using standard immunoassay testing.

#### Whole genome bisulfite sequencing data

Whole genome bisulfite sequencing (WGBS) data from obtained for 32 normal plasma and 26 hepatocellular carcinoma (HCC) patient plasma samples collected by Chan *et. al*. (24) was downloaded from the European Genome-phenome Archive (dataset accession: EGAS00001000566). An additional 4 HCC patient plasma and 4 normal plasma WGBS samples collected by Li *et. al*. (25) was downloaded from the European Genome-phenome Archive (dataset accession: EGAD00001004317).

### Mean locus methylation and weighted sample fraction of methylation density at the *ZNF154* locus from RRBS data

The RRBS data were aligned to hg19 using Bismark-0.19.0 (56). Counts of RRBS reads that overlapped the region Chr19:58220000-58220800 were tallied for each sample and divided into subgroups based on their specific start coordinates and combinations of methylated and unmethylated cytosines. Based on sample metadata, counts of reads from replicate libraries of the same sample were pooled together. One WBC sample did not have any reads at these coordinates and therefore was removed from the analysis.

Methylation density was defined as the number of methylated CpG dinucleotides (meCpGs) out of the total CpGs in a given read or DNA molecule. The weighted methylation level of a locus, or henceforth the mean locus methylation, was defined as the total number of meCpGs out of all CpGs sequenced in reads from the locus in question (57). The sample epiallelic fraction was defined as the proportion of reads with a given methylation density out of all reads from a given locus.

Reads with the same methylation density but having different numbers of CpGs were weighed in terms of their contribution to the overall sample methylation density by normalizing their epiallelic fractions in terms of the number of CpGs they covered out of the total CpGs covered in all of the reads. For this, the epiallelic fraction for a given set of reads was defined as:

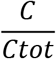

where *C* = # of CpGs covered by the reads with a methylation density above a given cutoff and *Ctot* = total CpGs in all reads.

### Methylation density classifier (EpiClass) procedure

The DREAMing assay includes two parameters that must be specified before a sample can be called positive: 1) a minimum methylation density MD_min_, and 2) a minimum epiallelic fraction EF_min_. A sample is considered positive if at least EF_min_ of the DNA fragments are methylated at a density above or equal to MD_min_, where the optimal parameters are chosen to maximize Youden J (58) (sensitivity + specificity, or, equivalently, true positive rate (TPR) – false positive rate (FPR)). Given a set of training data, EpiClass procedure solves for these two parameters simultaneously by calculating TPR-FPR for various combinations of MD_min_ and EF_min_, choosing the pair of parameter values that maximizes the value.

In practice, a range of values for MD_min_, were defined from 0 to 1 in increments of 0.05, with appropriate ranges of EF_min_ for each possible density MD_i_ determined by considering the full set of epiallelic fractions observed in the training data, at the selected MD_i_. In the extreme case that MD_min_=0, all fragments exhibiting a density above but not equal to MD_min_ were considered.

Epiallelic fractions for the RRBS data were calculated according to the weighted sample fraction method described above. For the plasma samples methylation density is derived from the melting temperatures as described below, and epiallelic fractions are calculated from the DREAMing melt peaks representative of each DNA divided by the total genomic equivalents assessed.

### Simulated dilution and classification performance comparison between the mean locus methylation and EpiClass using RRBS data

RRBS reads from EOCs or WBCs were randomly mixed together at different fractions at a set depth of total reads. A total depth of either 100, 1,000 or 10,000 reads was determined in order to simulate dilutions down to 0.01%. At each dilution, a simulated set of “spike-in” samples was made by randomly pairing each of the 12 EOCs with one of the 22 WBC samples for a total spike-in sample set size of 22. For each pair, a distribution of read methylation densities was generated for the EOC and the WBC sample based on the weighted sample fractions of methylation densities. Reads were randomly sampled from each distribution at a ratio equivalent to the % tumor dilution in question with the total number of reads sampled equal to the total read depth. For each dilution, receiver operating characteristic (ROC) curves were built for classifying samples using either the mean locus methylation level or the sample epiallelic fraction based on the methylation density cutoffs selected by EpiClass. The spike-ins were used as cases and the original 22 WBC samples as controls. ROC area under the curve (AUC) for the mean locus methylation and the optimal methylation density were recorded. This simulation was repeated 50 times and the mean AUC and the 95% CI for the mean locus methylation EpiClass methylation density were computed.

Using the same 50 iterations of the simulated dilution mentioned above, for select tumor dilutions (1.0%, 0.1%, 0.01%), AUC values were calculated for each methylation density cutoff as well as the sensitivity and specificity resulting from the optimal epiallelic fraction cutoff. The calculated AUCs were then used to determine the probability of achieving an improved performance over mean locus methylation for each methylation density cutoff. Given that the AUC can be defined as: “the probability that a randomly selected case will have a higher test result than a randomly selected control,” (59) each application of the methylation density cutoff was defined as a case and the resulting AUC was its test result, whereas the mean locus methylation AUCs would be the test results for a set of controls. New ROC curves based on these two sets of test results (i.e., sets of AUCs; n=50 for both methylation density [cases] and mean locus methylation [controls]) and the resulting AUC values of these ROC curves were used to calculate the probability that the methylation density cutoff would produce a higher AUC, or improved performance, than the mean locus methylation.

### Measurement of *ZNF154* methylation using DREAMing

#### Plasma cfDNA extraction and bisulfite conversion

cfDNA was extracted and processed according to the methylation-on-beads protocol (60) using NeoGeneStar Cell Free DNA Purification kits with pretreatment reagents and NeoGeneStar magnetic beads (NeoGeneStar, Somerset, NJ). Extracted cfDNA was bisulfite converted using Zymo Lightning Conversion reagents (Zymo Research, Irvine, CA) and eluted twice in 50 μl Zymo Elution Buffer using 1.5 ml LoBind Eppendorf tubes (Eppendorf, Hauppage, NY). Extracted cfDNA was quantified by qPCR in duplicate, using a primer and a TaqMan probe set to amplify a 100-bp region overlapping the bisulfite-converted top strand of the beta-actin locus on a C1000 Touch Thermo Cycler (BioRad, Hercules, CA) using a CFX96 Real-Time System (F: 5’-TAGGGAGTATATAGGTTGGGGAAGTT – 3’; R: 5’ – AACACACAATAACAAACACAAATTCAC – 3’; probe: /56-FAM/TGTGGGGTG/ZEN/GTGATGGAGGAGGTTTAG/3IABkFQ/).

#### Quantifying ZNF154 methylation in cfDNA with DREAMing

cfDNA *ZNF154* methylation densities were assessed by DREAMing according to previous established protocols (44). Briefly, based on the sample extraction yield, up to 4800 genomic equivalents from each cfDNA sample were distributed across 12 wells (approximately 400 equivalents per well) on a 96-well microtiter plate, for a total of 8 samples per plate. This was based on our ability to detect single fragments of fully methylated synthetic *ZNF154* target in a known quantity of low-methylation bisulfite-treated genomic DNA. As previously reported, the DREAMing method achieves single molecule quantification of the methylated cfDNA by using primers biased towards methylated DNA, assumes the methylated cfDNA fragments are rare in a background of cfDNA with low methylation density, and that the methylated cfDNA fragments are partitioned across the wells based on a Poissonian distribution. The unmethylated background cfDNA fragments all are expected to melt at the same temperature. However, wells that contain a methylated cfDNA fragment will produce a secondary melting peak whose melting temperature corresponds to the methylation density of the cfDNA fragment present in the well. Each sample was queried via DREAMing at least twice, for a total of 24 wells per sample, and the data for multiple runs of a given sample were pooled together. The fraction of methylated *ZNF154* genomic fragments for each sample was determined by counting the number of melting peaks above a defined temperature cutoff (corresponding to a specific methylation density cutoff, or number of methylated CpGs out of 14 total positions for our *ZNF154* locus; for conversion between these two values see next section), where each counted peak corresponded to a single methylated cfDNA fragment, inferred from Poissonian statistics.

The reaction conditions for DREAMing were as follows: Master PCR mixes were made so that each well would have a final volume of 25 μl with 200 μM dNTP mixture, 300 nM forward *ZNF154* DREAMing primer (5’ – GGGCGATATTGGTAGGGATT – 3’), 300 nM reverse *ZNF154* DREAMing primer (5’ – AAATATATTCACCGAATCAAAAATAACAAAA – 3’), 1X EvaGreen (Biotium Inc, Fremont, CA), 0.04 U/μl Platinum Taq (Thermo Fisher Scientific, Bothell, WA), and 1X in-house “Magic Buffer” (16.6 mM ammonium sulfate, 67 mM Tris pH 8.8, 2.7 mM magnesium chloride, and 10 mM beta-mercaptoethanol). DREAMing reactions were run on a C1000 Touch Thermal Cycler using a CFX96 Real-Time System. Reactions were run at 95°C for 5 minutes for 1 cycle; 95°C for 30 seconds, 61.4°C for 30 seconds, and 72°C for 30 seconds for 50 cycles; followed by a temperature gradient beginning at 65°C and ramping up to 90°C in 0.2°C increments, each held for 10 seconds, before SYBR/FAM fluorescence was imaged. After DREAMing, melting temperature peaks were visualized using the accompanying CFX Manager 3.1 software to analyze the negative derivative of the change in fluorescence (-d(RFU)/dT) versus temperature plots for each well.

#### Sample epiallelic fractions

The volume of sample plasma assed in DREAMing was determined by the amount of genomic equivalents (derived from beta-actin qPCR measured copies) loaded into the assay for a given sample divided by the total number of beta-actin copies measured in the entire sample elution, multiplied by the starting volume of sample plasma processed. The counts of epialleles of a given methylation density measured by DREAMing were then divided by this number to return: counts of epialleles per mLs of plasma.

#### Validation and calibration of DREAMing melt peak temperatures to methylation densities

Bisulfite amplicon sequencing was used to validate the results of at least one DREAMing well per sample in the training set. Wells exhibiting a temperature peak indicative of a high methylation density cfDNA fragment were preferentially selected and, when possible, multiple wells representative of the overall methylation profile were selected for each respective sample.

Sequencing was performed by pipetting 20 μl from chosen wells into a separate 96-well plate and cleaned using Ampure XP beads (Beckman Coulter, Brea, CA) according to the manufacturer’s protocol, at a ratio of 1.8 μl beads to 1 μl sample. The DNA was then eluted with 35 μl EB buffer (Qiagen, Germantown, MD), and 30.75 μl of the elution was combined with 5 μl 10X TaKaRa EpiTaq PCR Buffer (Mg^2+^ free; TaKaRa, Mountain View, CA), 5 μl 25 mM MgCl_2_, 6 μl 2.5 mM dNTP mixture, 1 μl 12.5 μM forward primer (175-bp forward *ZNF154* DREAMing primer with sequencing adapter), 1 μl 12.5 μM reverse primer (175-bp reverse *ZNF154* DREAMing primer with sequencing adapter), 1 μl DMSO, and 0.25 μl of 5 U/μl TaKaRa EpiTaq DNA polymerase, for a total reaction volume of 50 μl. This mixture was placed in a SimpliAmp thermal cycler (Applied Biosystems, Foster City, CA) using the following conditions: 95°C for five minutes and one cycle; 95°C for 30 seconds, 50°C for 30 seconds, and 72°C for 30 seconds, for nine cycles; and 72°C for seven minutes for one cycle. A second Ampure XP beads cleanup was performed by combining 46 μl each PCR reaction with 55 μl beads, eluting in 27 μl EB buffer. Next, a 23 μl elution was combined with 25 μl 2X High Fidelity Phusion Master Mix (NEB, Ipswich, MA), and 1 μl i7 and i5 barcoding primers for a total reaction volume of 50 μl. Another round of PCR was performed under the following conditions: 98°C for 30 seconds and one cycle; 98°C for 10 seconds, 65°C for 30 seconds, and 72°C for 30 seconds, for nine cycles; and 72°C for five minutes and one cycle. After this, each reaction was cleaned again with Ampure XP beads using 55 μl beads and 46 μl sample and eluted in 30 μl EB buffer. Then a 3 μl elution was run on a 2% agarose gel to confirm the expected band of 300-bp (size of amplicon with adapter and barcodes). Samples were submitted to the NIH Intramural Sequencing Center for quality control and sequencing on a MiSeq using 300-bp paired end sequencing. Using Bismark-0.19.0, analysis of reads, bisulfite conversion efficiency, and determination of meCpG patterns was performed as described previously (41). Wells with sequenced amplicons that had less than 95% bisulfite conversion efficiency were discarded.

DREAMing melt peak temperatures were converted to methylation density values by linear regression of methylation density values determined by the bisulfite amplicon sequencing. Briefly, sequencing patterns were ordered based on methylation density and abundance and the pattern with the highest methylation density from this list was then matched to the highest melt peak temperature, the results of which are shown in Supplementary Fig S7. The generated linear model was then used to convert all melt peak temperatures into methylation densities and rounded these to the nearest 7% given that we would expect each additional meCpG in our locus of 14 potential meCpG sites to increase the methylation density by 1/14 or approximately 7%.

#### Mean locus methylation in plasma

The epiallelic fraction of all detected epialleles via DREAMing except fully unmethylated DNA fragments (based on a methylation density cutoff of 0%) was used as an estimate of the sample mean locus methylation. This was based on the fact that, unlike the RRBS reads, the number of CpG sites covered (14) was the same for each DNA fragment targeted in the DREAMing assay. Therefore, the fraction of these CpGs that were methylated would be proportional to the fraction of total methylated epialleles.

### Comparison to CancerDetector

#### Overview

The combined 30 HCC and 36 normal WGBS plasma samples were randomly split 50:50 ten times into training and test sample set pairs, where each pair was considered a separate run. Training samples in each run were used by EpiClass or CancerDetector to define an optimal methylation density cutoff (with respect to EpiClass) or Tumor Class and Normal Class beta distribution shape parameters, for each marker (see Li. et al. (25) for detailed description). The methylation density cutoffs were applied to the test sample sets to construct ROC curves based on sample counts of methylated reads at or above the cutoff in order to assess the performance on EpiClass, or the Class shape parameters were used to estimate tumor fractions in the test set samples, which were used to construct ROC curves in order to assess the performance of CancerDetector.

#### Preprocessing of WGBS datasets

The WGBS data were aligned to hg19 using Bismark-0.20.0 (56). Reads adapters were trimmed with TrimGalore-0.6.0 (61) and duplicate reads were removed with `deduplicate_bismark. bismark_methylation_extractor` was used to generate coverage files indicating the number of unmethylated and methylated CpGs in the aligned reads at each CpG position. For each marker, reads that were aligned to the genomic region of interest were extracted. Reads which did not overlap completely were ignored.

#### EpiClass pipeline

Read methylation density tables of the WGBS samples for each marker region were generated using the `epiclass READtoMD` command. For each training and test sample set pair, the optimal methylation density cutoff for normalized read counts was determined using the `epiclass MDBC` command on the training set, then run again on the test set with the flag `sampleValsAtMD` to obtain sample read counts with methylation densities at or above the methylation density cutoff.

#### CancerDetector pipeline

For each run, Tumor Class and Normal Class beta distribution shape parameters were determined for each marker, as described in Li. et al. (25). In brief, BMIQ-normalized LIHC 450K methylation array data was filtered for probes within the marker region for 50 tumor and normal sample pairs. Probes overlapping SNPs were removed. Beta values at each of the probes for the tumors were used to learn the shape parameters for the Tumor Class beta distribution and conversely beta values at each of the probes for the normal samples were used to learn the shape parameters for the Normal Class beta distribution. In addition, the bismark CpG coverage files derived from `bismark2bedGraph` were used to determine the methylation level at each CpG site within the given marker region for the normal plasma WGBS samples and these values were added to the Normal Class beta distribution for each corresponding training set. Once shape parameters were learnt, these were recorded and used as inputs into CancerDetector. Methylation patterns of reads overlapping each marker region of interest were also used as input for test set samples to determine their tumor fraction values.

#### Marker regions

For *ZNF154*, we used the marker region chr19:58220195-58220937, which was used previously as a marker region in CancerDetector for liver cancer (25). To identify additional markers hypermethylated in liver cancer, we computed the median difference in beta values using TCGA LIHC 450K methylation array data for CpG probes between the 50 matched tumor and normal LIHC samples and took the top 3 that i) did not correspond to a SNP, ii) the maximum beta value in normals was < 0.2, iii) were hypermethylated in tumors relative to normals, iv) overlapped with a marker region previously defined in the CancerDetector study. This resulted in: marker 6263 chr2:208989109-208989679; marker 5594 chr2:127782982-127783470; marker 29305 chr13:107187077-107187512.

#### EpiClass combined marker analysis

When classifying test set samples using multiple markers, a support vector machine classifier using a linear kernel was trained on the normalized methylated read counts above the methylation density cutoff for each marker region being assessed for each sample of the training set. The classifier was then applied to the test sample set. This was done using sklearn.svm.SVC(kernel=‘linear’, probability=True, random_state=0) from sklearn (v0.23.1) (62).

Lists of training and test sample set shape parameters used in CancerDetector and the EpiClass optimal methylation density cutoffs can be found in Supplementary Table S3.

### Statistical analyses and plotting

Plotting and statistical analyses were performed using custom Python-3.7.4 (https://python.org) and R-3.4.4 (63) scripts available at https://github.com/Elnitskilab/EpiClass. To compare epiallelic fractions or mean locus methylation between groups, boxplot comparisons were performed. Statistical significance was evaluated using the two-sided Wilcoxon rank sum test. The true and false positive rates, associated thresholds, as well as the area under the curve (AUC) for the receiver operating characteristic analyses were generated using the python package sklearn (v0.23.1) (62) with the module sklearn.metrics. AUC 95% confidence intervals were computed using the R library pROC (64) and using the ci.auc() command with method=”bootstrap” on the data using 2000 stratified replicates. Statistical significant difference between ROC curves was computed using the roc.test() command.

## Abbreviations

HRM: high resolution melt
DREAMing: Discrimination of Rare EpiAlleles by Melt
ctDNA: circulating tumor DNA
WGBS: whole-genome bisulfite sequencing
cfDNA: cell-free DNA
EpiClass: methylation density classifier
MDC: methylation density cutoff
RRBS: reduced representation bisulfite sequencing
EOCs: epithelial ovarian carcinomas
CGI: CpG island
meCpGs: methylated CpG dinucleotides
TPR: true positive rate
FPR: false positive rate
ROC: receiver operating characteristic
AUC: area under the curve
HCC: hepatocellular carcinoma
TCGA: The Cancer Genome Atlas
LIHC: liver hepatocellular carcinoma

## Declarations

### Ethics approval and consent to participate

The study was conducted in accordance with the Declaration of Helsinki, was approved by the Institutional Review Board at the National Cancer Institute and registered with Clinicaltrials.gov (NCT01445509 and NCT00436215). All patients provided written informed consent at study enrollment.

### Consent for publications

Not applicable.

### Availability of data and materials

*450K Illumina Infinium HumanMethylation450 BeadChip datasets.* Processed Illumina Human 450K data for EOCs (GEO accession GSE72021), and WBCs (GEO accession GSE55763) was downloaded from the NCBI’s Gene Expression Omnibus (GEO, https://www.ncbi.nlm.nih.gov/geo/). Data for TCGA solid tumor and control sample sets was downloaded from the Broad Institute (https://gdac.broadinstitute.org/) FireHose.

*Reduced representation bisulfite sequencing data.* The data was downloaded from the European Genome-phenome Archive (dataset accession: EGAD00001003822).

*Whole genome bisulfite sequencing data.* The data was downloaded from the European Genome-phenome Archive (dataset accessions: EGAS00001000566, EGAD00001004317).

The EpiClass Python code and input files are available at the GitHub website: https://github.com/Elnitskilab/EpiClass. This GitHub project is intended to demonstrate the steps to generate the analyses and figures presented in this study. Work is underway to extend the program for general use with other datasets and genomic loci. Interested parties may contact the authors for more details. Here, too, code to reproduce figures in this study or describing the CancerDetector comparison are available. The CancerDetector code itself should be requested from Li *et. al*. (25).

### Preprint sharing and citation

A preprint of this work has been deposited into the bioRxiv preprint server: https://doi.org/10.1101/579839.

### Authors’ contributions

BFM, TRP, and LE conceptualized the manuscript. BFM, TRP, HP, and PA performed the experiments. BFM and GM analyzed and visualized the data. BFM and TRP wrote the original draft. BFM, TRP, and LE reviewed and edited the manuscript. LE, TRP, TW, and CMA provided funding and support. BFM and AG wrote and packaged the executable code. LE and TRP supervised the project. AO and CMA provided samples.

## Acknowledgements

We would like to thank Kristin Harper and Karoun Bagamian for helpful edits and useful feedback during the writing of this piece, Ryan Winters for supplying us with plasma samples from the Fox Chase Cancer Biorepository, Alice Young for guidance during sequencing library preparation and data acquisition, Dr. Martin Widschwendter for access to the RRBS datasets used in this study, Dr. Jasmine Zhou for access to the WGBS datasets used in this study, Dr. Wenyuan Li for access to the CancerDetector program, Alejandro Stark for instruction on cfDNA extraction and processing, and Thor Nilsen for technical assistance with the NeoGeneStar cfDNA Purification Kit.

---

**Supplementary Figure S1:**
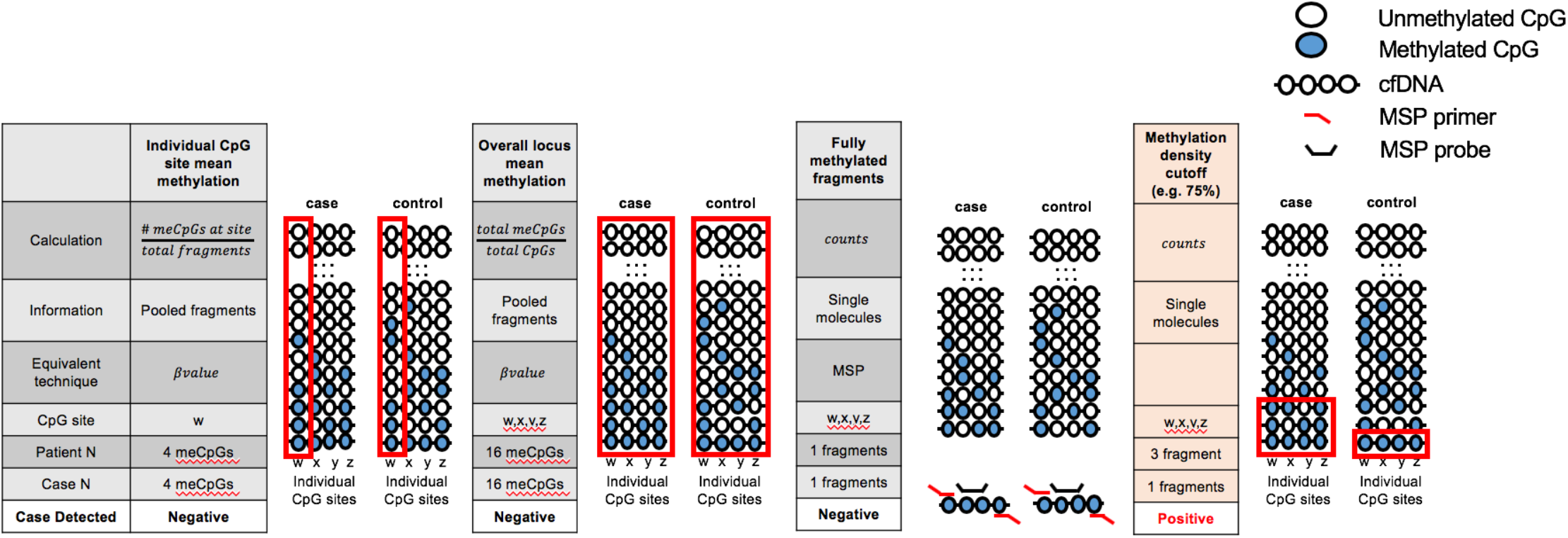
Illustration of different approaches for calculating methylation differences between case and control cfDNA epialleles. Case and control are the same examples from Figure 1. Red outlines indicate the read methylation information that is assessed by each approach. Fully-methylated fragments representative of a target queried in methylation-sensitive PCR (MSP) assays. Here, both case and control have equal counts of fully methylated molecules for which the probe can anneal to the target cfDNA (after bisulfite conversion) and are indistinguishable.

**Supplementary Figure S2:**
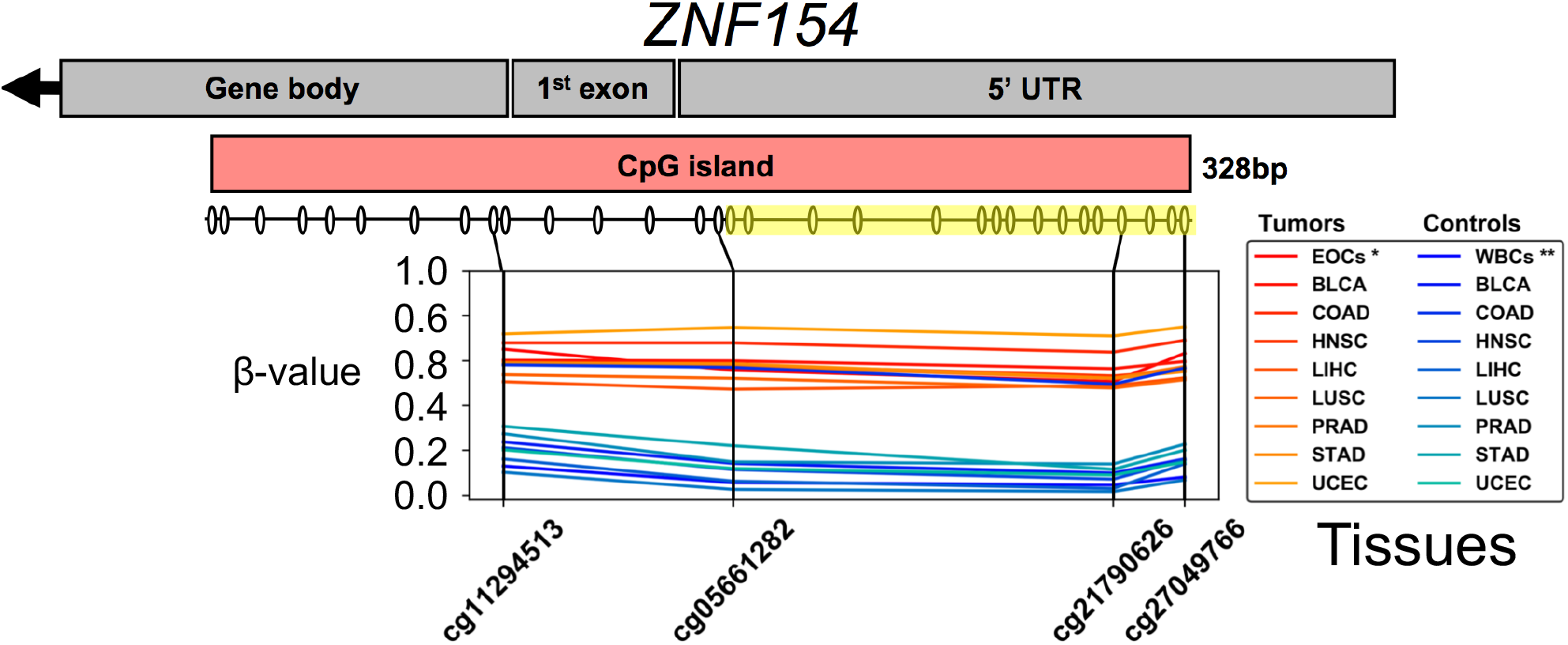
Methylation at the *ZNF154* genomic locus. **A**) The *ZNF154* gene is encoded on the reverse strand of Chromosome 19 and contains a 328-bp CpG island (CGI) that extends from the 5’-UTR through into the *ZNF154* gene body itself. Schematic showing β-values at multiple CpG sites determined from Illumina Infinium HumanMethylation450 array data for tumor (*red lines*) and control (*blue lines*) tissues. The target locus assessed in the DREAMing assay is highlighted (*yellow*). Abbreviations: ovals below the CpG island represent CpG positions; EOCs = epithelial ovarian carcinomas (n=221); WBCs = white blood cells; * indicates data taken from Widschwendter *et. al* (8); ** indicates data from Lehne *et. al* (46); remaining 4 letter acronyms correspond to TCGA tissue codes: BLCA = bladder carcinoma (tumors = 412, controls = 21); COAD = colon adenocarcinoma (tumors = 295, controls = 38); HNSC = head-neck squamous cell carcinoma (tumors = 528, controls = 50); LIHC = liver hepatocellular carcinoma (tumors = 377, controls = 50); LUSC = lung squamous cell carcinoma (tumors = 370, controls = 42); PRAD = prostate adenocarcinoma (tumors = 498, controls = 50); STAD = stomach adenocarcinoma (tumors = 395, controls = 2); UCEC = uterine corpus endometrial carcinoma (tumors = 431, controls = 46).

**Supplementary Figure S3:**
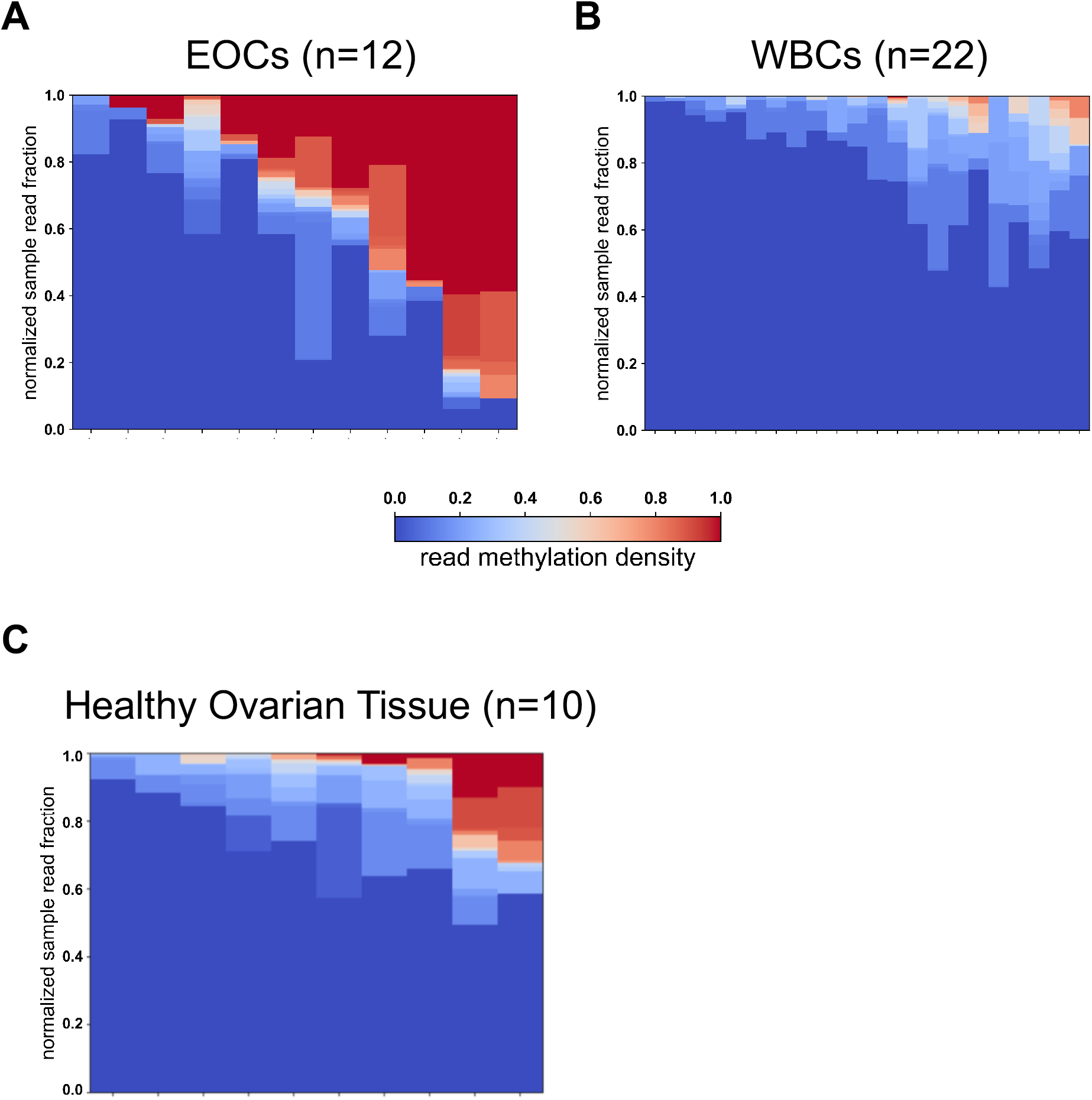
Methylation density at the *ZNF154* genomic locus. **A-C**) Heatmaps showing the relative fractions and corresponding methylation density profiles derived from RRBS reads of the *ZNF154* target locus for ovarian carcinomas (n=12), healthy ovarian tissues (n=10), and WBCs (n=22). Abbreviations: WBCs = white blood cells. Data from Widschwendter *et. al* (8).

**Supplementary Figure S4:**
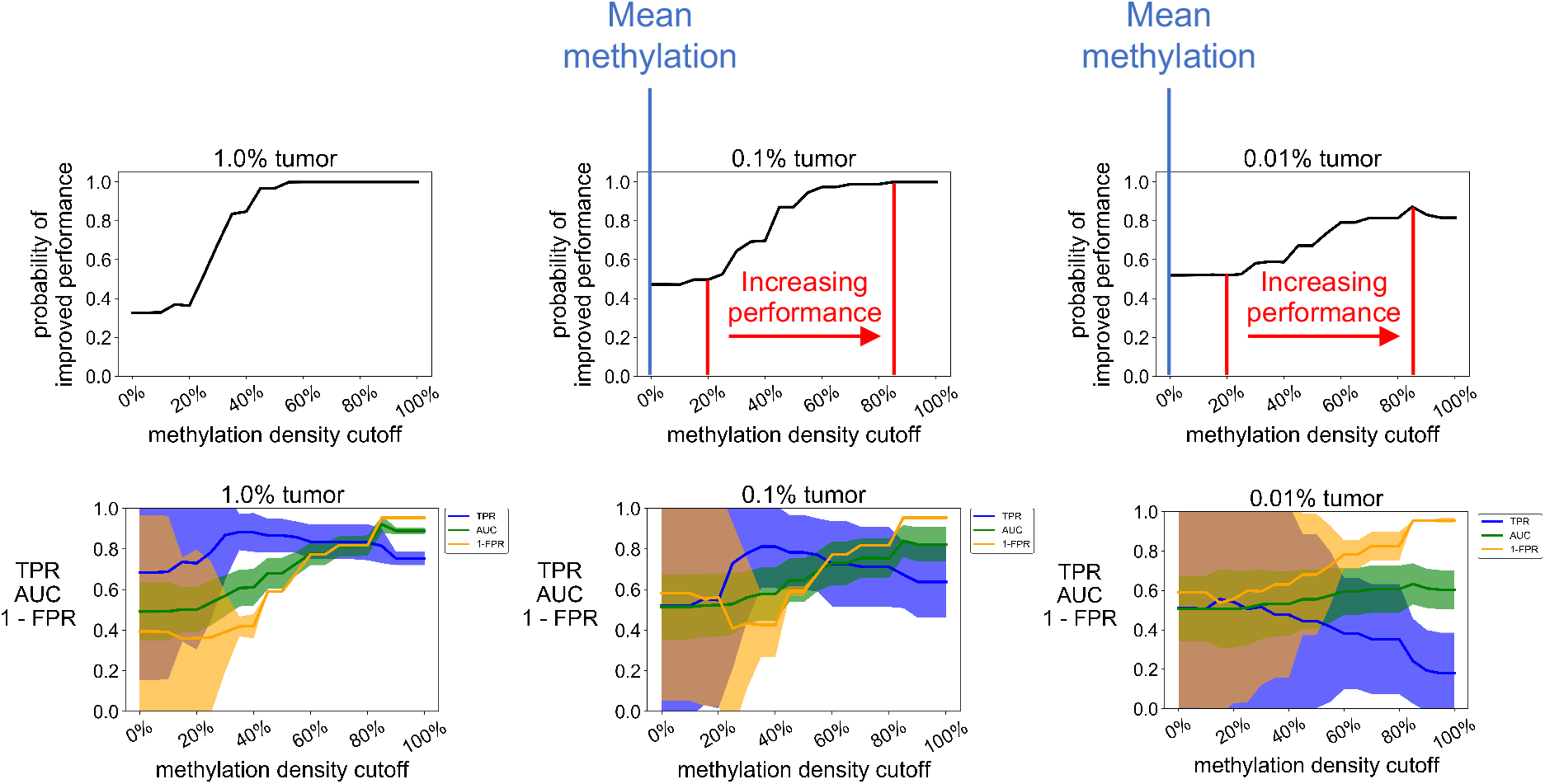
Simulated performance of EpiClass using varying admixture ratios of ovarian carcinoma (EOC) to WBC RRBS reads at 10000 reads per sample. Plots showing the probability of achieving a higher AUC using EpiClass compared to the mean locus methylation classifier for each methylation density cutoff at various admixture ratios. For sub-1% tumor fractions, the range of methylation density cutoffs (20%-85%) that result in increasing probability of improved classification performance over mean methylation is indicated between red lines. Mean methylation methylation density cutoff indicated by blue lines. Lower panels show the TPR, 1-FPR, and AUC achieved by using EpiClass at each methylation density cutoff. Solid lines indicate the mean value and shaded regions indicate the 95% confidence interval for 50 iterations of the simulation. Abbreviations: AUC = area under the curve; TPR = true positive rate; FPR = false positive rate.

**Supplementary Figure S5:**
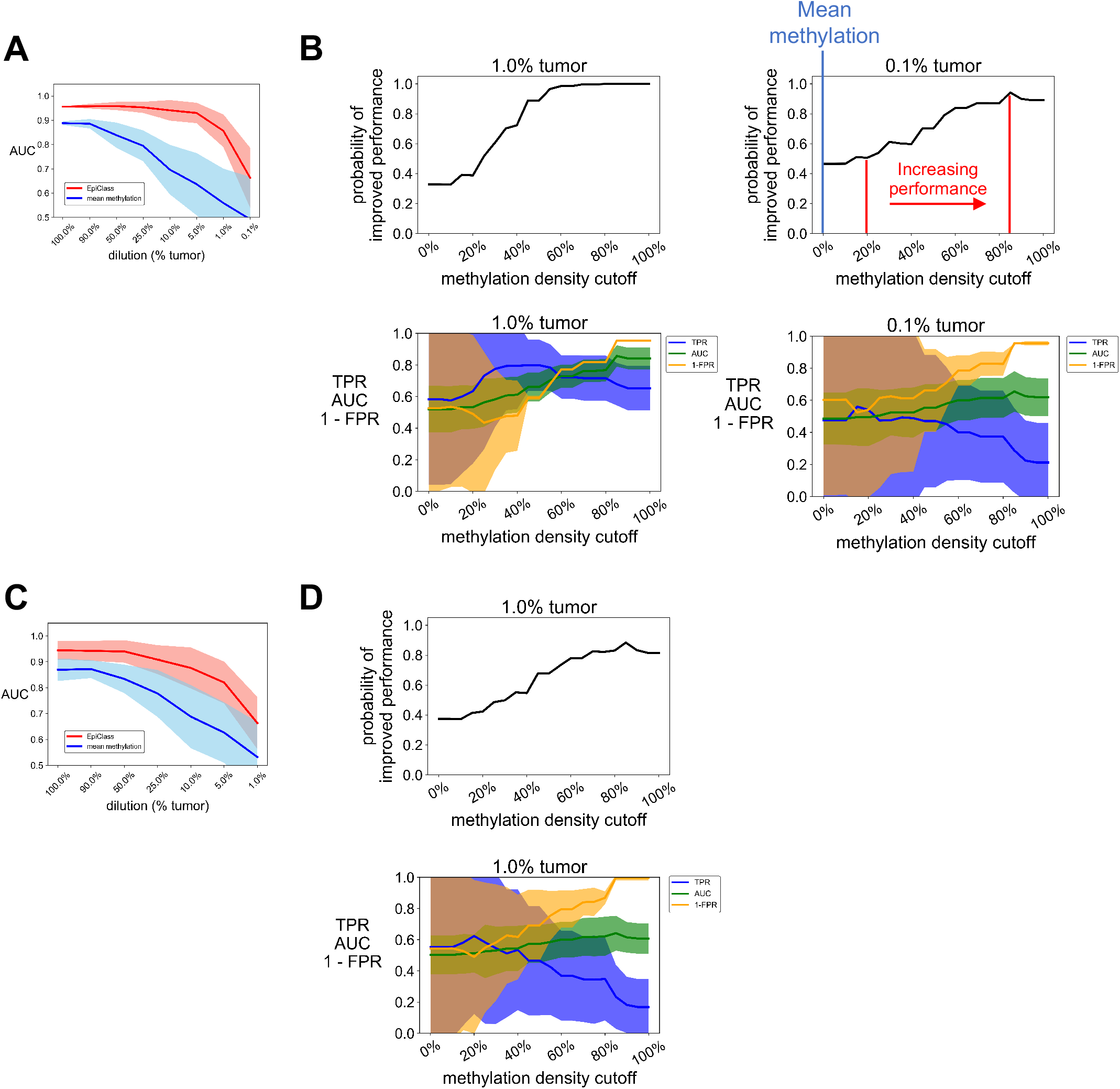
Simulated performance of EpiClass using varying admixture ratios of ovarian carcinoma (EOC) to WBC RRBS reads sampled at 100 and 1000 total reads per simulated sample. **A**) The performance of the methylation density binary classifier (EpiClass, *red*) and mean locus methylation classifier (*blue*) at increasing dilutions of EOC RRBS reads in a background of WBC RRBS reads acquired from Widschwendter *et al*. (8) with 1000 total reads sampled. **B**) Plots showing the probability of achieving a higher AUC than the mean locus methylation classifier using EpiClass for each methylation density cutoff at various admixture ratios based on 1000 read per simulated sample. Lower panels show the TPR, 1-FPR, and AUC achieved by using EpiClass at each methylation density cutoff. EOCs (n=12) were randomly paired with a WBC (n=22) sample and RRBS reads were sampled from an EOC-WBC pair to generate a simulated spike-in sample. Simulated samples (n=12) were compared to the original WBC samples (n=22). 50 iterations of the simulation were performed. Solid lines indicate mean values and shaded regions indicate 95% confidence intervals. For sub-1% tumor fractions, the range of methylation density cutoffs (20%-85%) that result in increasing probability of improved classification performance over mean methylation is indicated between red lines. Mean methylation methylation density cutoff indicated by blue lines. **C-D**) Same as A) and B) except with 100 total reads sampled. Abbreviations: EpiClass = methylation density classifier; AUC = area under the curve; TPR = true positive rate; FPR = false positive rate.

**Supplementary Figure S6:**
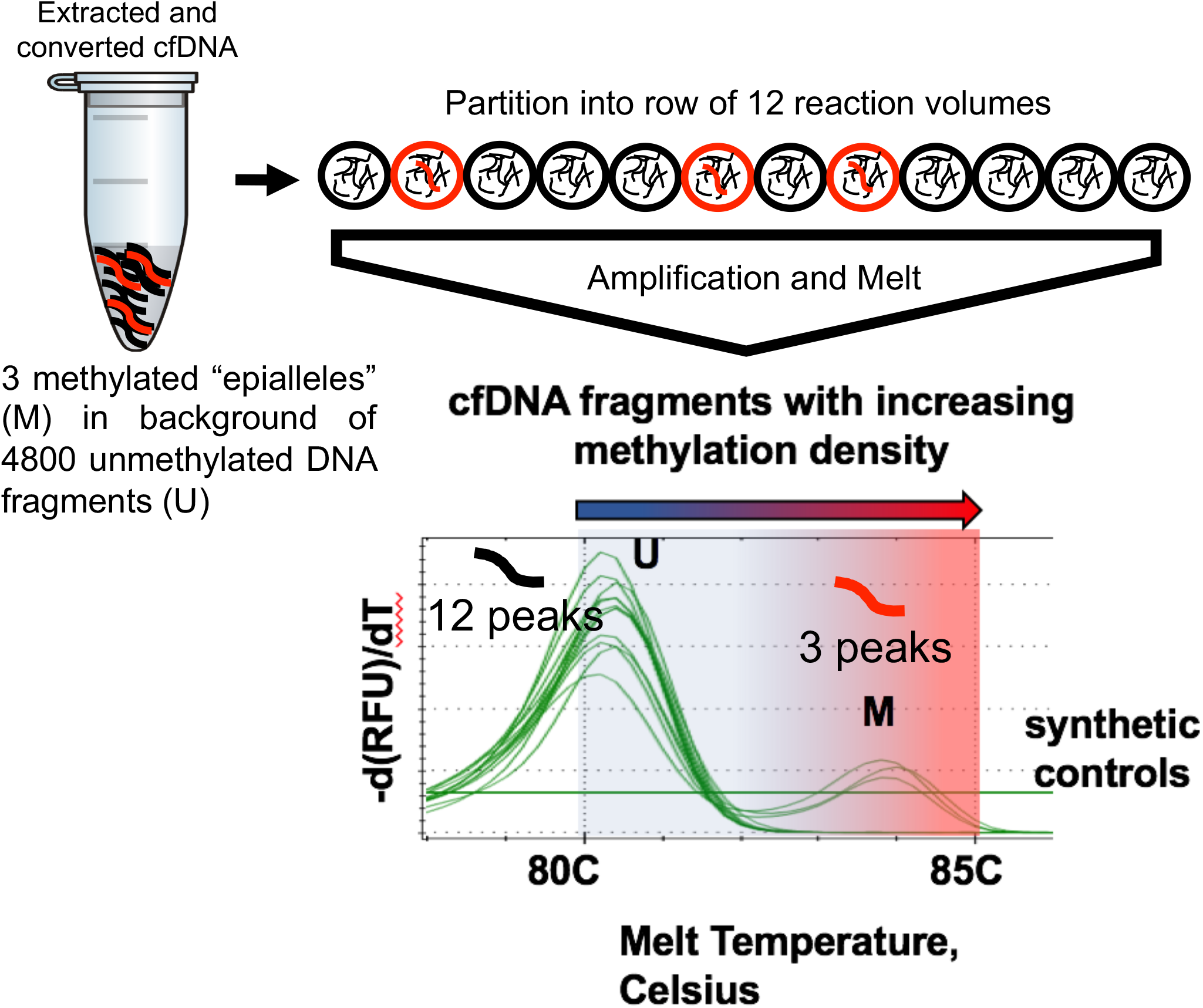
Melting traces from DREAMing assay for 12 wells containing in total a mixture of 3 synthetic fully-methylated (M, red) and 4800 unmethylated control male genomic DNA (U, black) fragments (400 U per well) originating from the ZNF154 genomic region of interest. The sample is partitioned such that there is no more than 1 methylated epiallele per well (in addition to unmethylated DNA). The 3 rare methylated epialleles are assumed to be distributed among the 12 wells based on a Poissonian distribution. All wells give a melt peak indicative of the presence of unmethylated U background DNA fragments, which all melt at approximately the same temperature. Wells with a methylated epiallele produce a secondary melt peak corresponding to the methylation density of the epiallele. Abbreviations: -d(RFU)/dT = negative derivative of the change in relative fluorescent units. U = unmethylated DNA; M = methylated DNA.

**Supplementary Figure S7:**
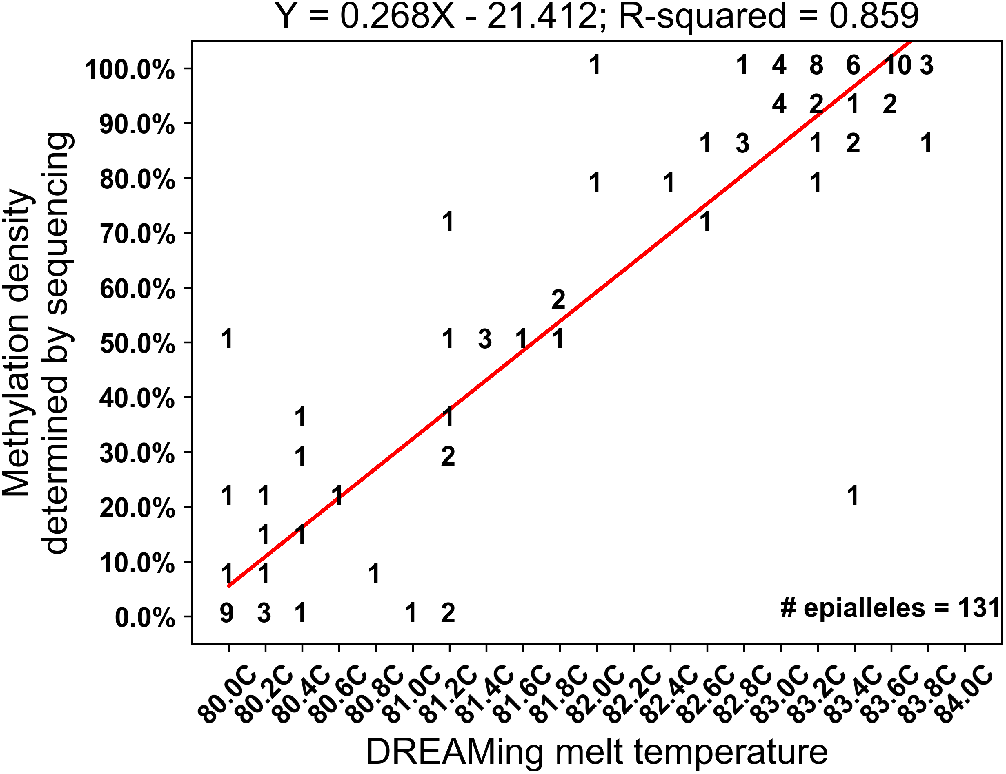
*ZNF154* locus methylation density vs. DREAMing melt temperature in cfDNA. DREAMing melt peak temperatures and corresponding methylation density measurements identified via bisulfite sequencing for 131 post-DREAMing epiallele amplicons. The numbers on the plot represent the number of times a detected melt peak had the corresponding melt temperature and produced an amplicon with the corresponding methylation density. The red line denotes the best fit regression line. The linear regression model and R-squared value is shown above the plot.

**Supplementary Figure S8:**
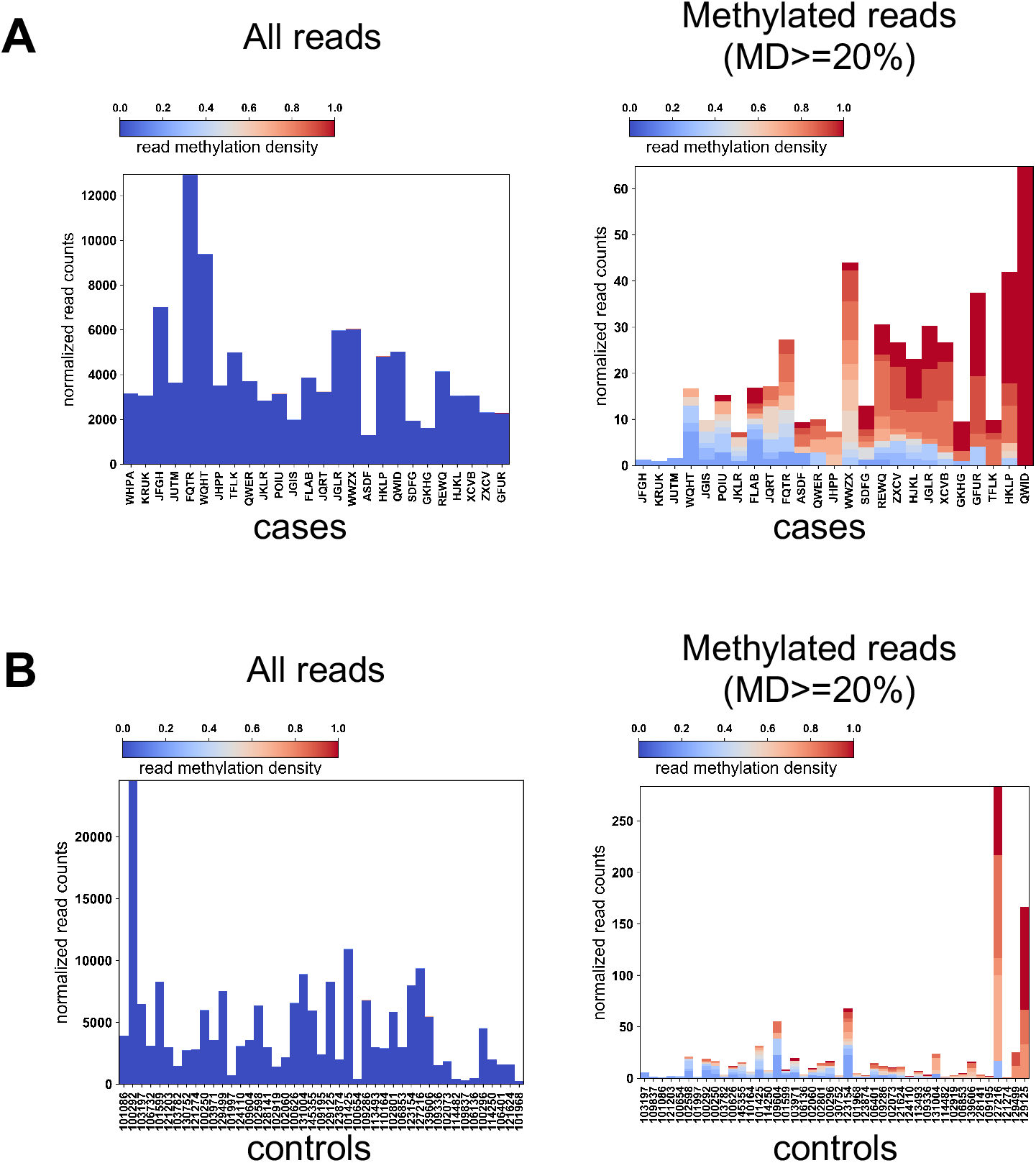
Fraction of total reads and methylated reads in case (A) and control (B) samples from the training cohort. Methylated reads are considered epialleles with a methylation density >= 20%. Y-axes indicate the normalized counts of reads (epialleles per mL that were quantified in DREAMing) for each sample. Color bars indicate the methylation density of the quantified reads. MD = “methylation density”. X-axes indicate sample IDs.

**Supplementary Figure S9:**
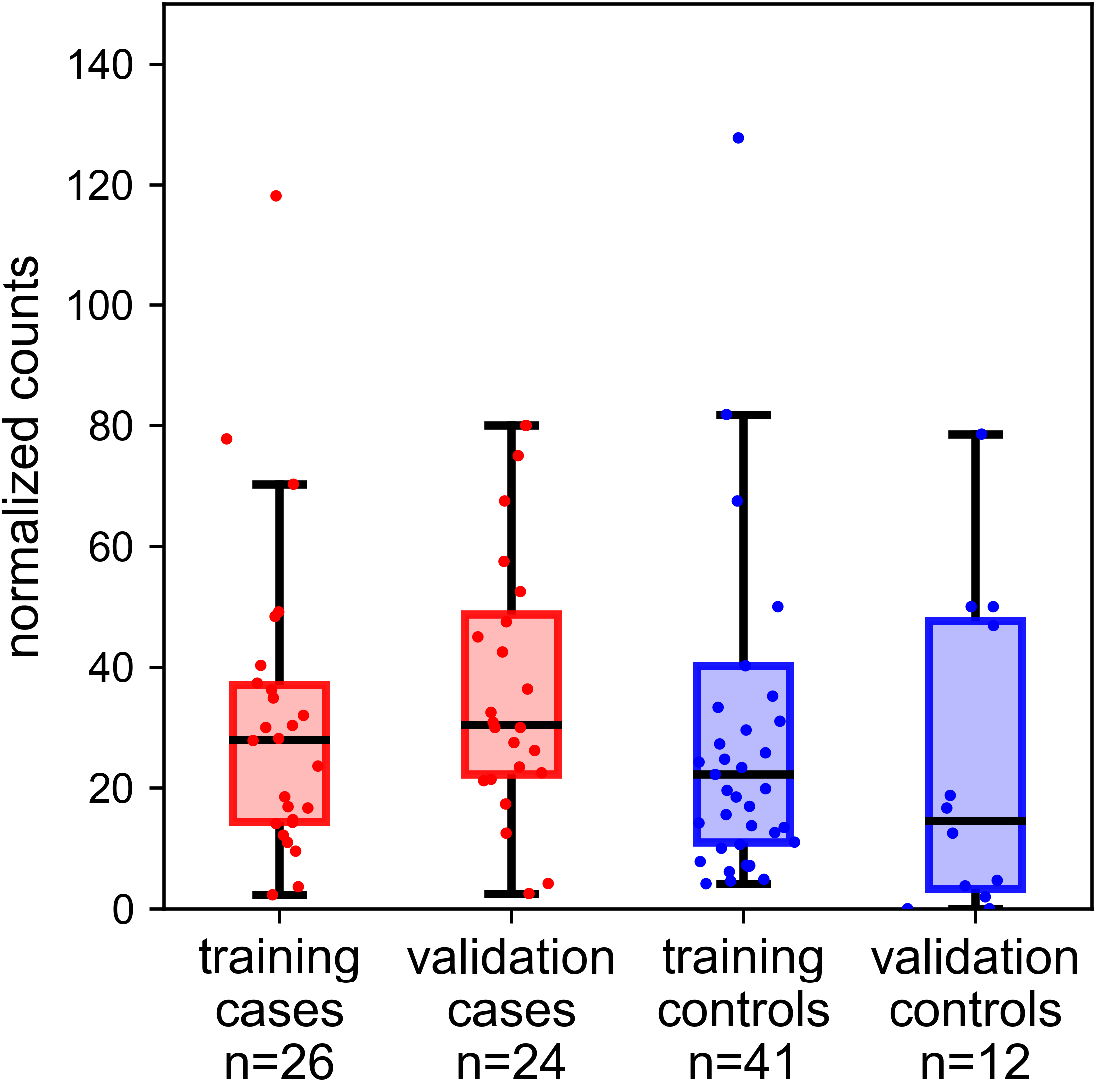
Total normalized counts (counts per mL sample plasma) of methylated reads with any methylation in cases and controls from the training and validation sample cohorts. Methylated reads include all reads with a methylation density > 0%. No statistical difference between the sample cohorts; two-sided Wilcoxon rank-sum test: Training cases median counts = 28.0; Validation cases median counts = 30.4; Training controls median = 22.2; Validation controls median = 14.6; rank sum p-values: Training vs. Validation cases p = 0.236; Training vs. Validation controls p = 0.126; Training cases vs. controls p = 0.719; Validation cases vs controls p = 0.058.

**Supplementary Figure S10:**
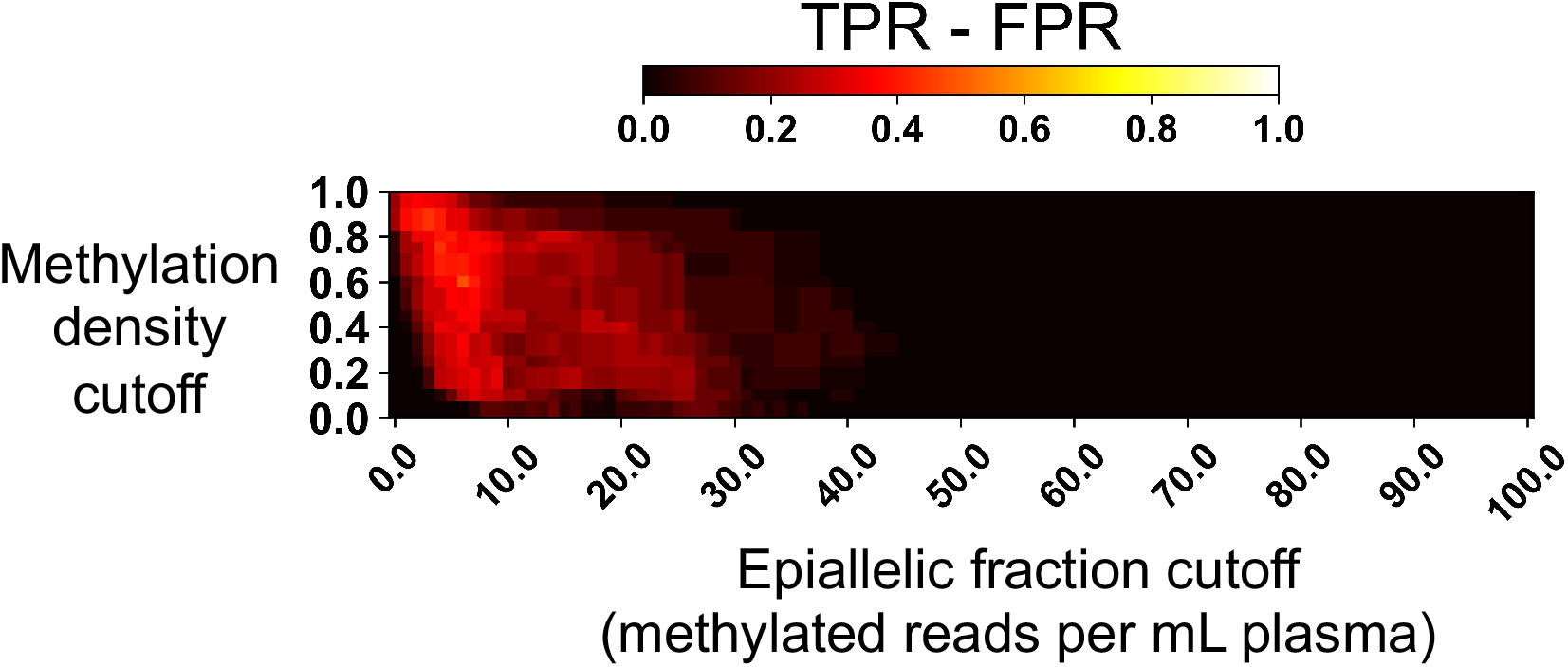
EpiClass heatmap indicating the true and false positive rate differences for each combination of methylation density and epiallelic fraction cutoffs for identification of EOC (n=26) versus healthy control (n=41) plasma samples.

**Supplementary Figure S11:**
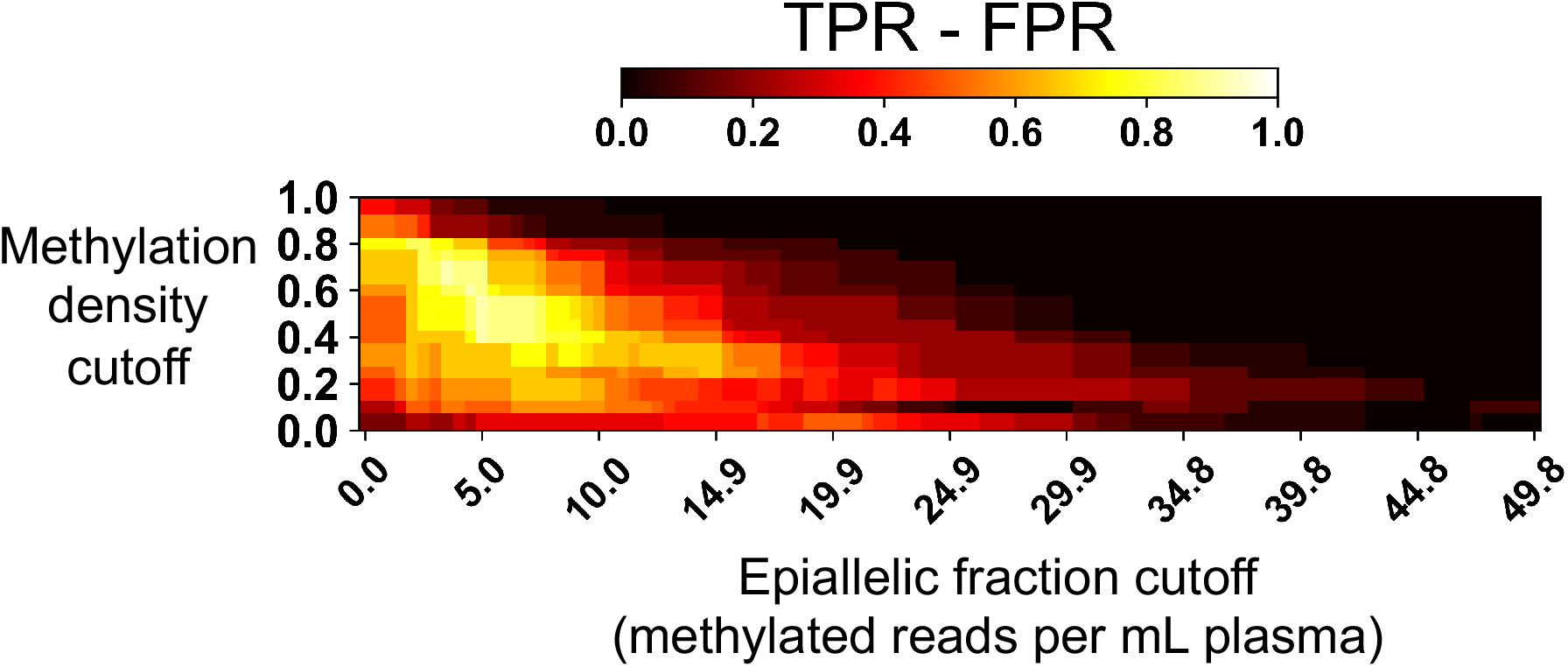
EpiClass analysis of cfDNA DREAMing data from the validation cohort. **(A)**EpiClass heatmap indicating the true and false positive rate differences for each combination of methylation density and epiallelic fraction cutoffs for identification of EOC patient (n=24) versus healthy control (n=12) plasma samples of the second cohort.

